# *Arabidopsis* cell surface LRR immune receptor signaling through the EDS1-PAD4-ADR1 node

**DOI:** 10.1101/2020.11.23.391516

**Authors:** Rory N. Pruitt, Lisha Zhang, Svenja C. Saile, Darya Karelina, Katja Fröhlich, Wei-Lin Wan, Shaofei Rao, Andrea A. Gust, Federica Locci, Matthieu H.A.J. Joosten, Bart P.H.J. Thomma, Jian-Min Zhou, Jeffery L. Dangl, Detlef Weigel, Jane E. Parker, Farid El Kasmi, Thorsten Nürnberger

## Abstract

Plants use both cell surface and intracellular immune receptors with leucine rich-repeat (LRRs) to detect pathogens. LRR receptor kinases (LRR-RKs) and LRR receptor-like proteins (LRR-RPs) recognize extracellular microbe-derived molecules to confer pattern-triggered immunity (PTI), while nucleotide-binding LRR (NLR) proteins detect microbial effectors inside the cell to confer effector-triggered immunity (ETI). Despite PTI and ETI signaling being initiated in different compartments, both rely on the transcriptional activation of similar sets of genes, suggesting convergence in signaling upstream of nuclear events. Here we report that two sets of molecules, helper NLRs from the ADR1 (ACTIVATED DISEASE RESISTANCE 1) family as well as lipase-like proteins EDS1 (ENHANCED DISEASE SUSCEPTIBILITY 1) and PAD4 (PHYTOALEXIN DEFICIENT 4), are required not only for ETI, but also for PTI. A further similarity is seen in the evolutionary patterns of some PTI and ETI receptor genes, with both often being highly polymorphic, and with nevertheless distinct roles of LRR-RK and LRR-RP receptors in immunity. We find that the LRR-RK SOBIR1 directly links LRR-RPs with the ADR1 helper NLR as well as EDS1 and PAD4, suggesting the formation of constitutive supramolecular signalosome complexes at the inner side of the plasma membrane. We propose that the EDS1-PAD4-ADR1 node is an essential component and convergence point for immune signaling cascades activated by both surface-resident LRR-RP receptors and intracellular NLR receptors.

## Introduction

Plants employ a two-tiered immune system to combat microbial invasion^1^. Cell surface pattern recognition receptors (PRRs) recognize conserved microbial surface structures (pathogen-associated molecular patterns, PAMPs) to elicit pattern-triggered immunity (PTI). PTI confers full resistance to host non-adapted pathogens and partial (basal) resistance to host-adapted pathogens. Leucine rich-repeat receptor kinases (LRR-RKs) and receptor-like proteins (LRR-RPs) are two major classes of plant PRRs that confer PTI through sensing proteinaceous microbial patterns. In contrast to LRR-RKs, LRR-RPs lack a cytoplasmic protein kinase domain and form constitutive heteromeric complexes with the LRR-RK SOBIR1^2,3^. Upon ligand binding, LRR-RKs and LRR-RP/SOBIR1 complexes recruit the LRR-RK BAK1 to initiate intracellular signal transduction and immune activation^3–6^. The contrasting roles of the *Arabidopsis thaliana* (hereafter *Arabidopsis*) receptor-like cytoplasmic kinase (RLCK) BIK1 (BOTRYTIS-INDUCED KINASE 1) in LRR-RK and LRR-RP PTI activation, as well as quantitative and qualitative differences in downstream defense responses triggered by either PRR type, suggest immune receptor type-dependent diversification of signaling pathways in PTI^7^.

Intracellular nucleotide-binding domain leucine rich-repeat (NLR) immune receptors recognize strain-specific microbial effectors to activate effector-triggered immunity (ETI) and host cell death^8^. NLRs are classified as coiled-coil (CC), TOLL-INTERLEUKIN 1 RECEPTOR (TIR) or RPW8-like CC (CCR; HELO-domain) NLRs based on the different structures of their N-terminal domains^9^. Two subfamilies of CCR-type NLRs, with ADR1 and NRG1 as founding members, function downstream of many sensor CC-NLRs and most, if not all, TIR-NLRs and are thus considered as helper NLRs (hNLRs)^10–12^. In *Arabidopsis*, the two hNLR groups function together with the EDS1-family of lipase-like proteins to relay signals downstream of sensor NLRs in ETI^10,13–15^. While genes from the ADR1 and NRG1 families seem to be conserved between individuals of the same species, genes encoding NLRs are notable for their pronounced presence/absence patterns, often being present only at intermediate frequencies^16,17^. Defense outputs induced upon activation of PTI or ETI are qualitatively similar, but whether and to what extent signaling pathways underlying immune activation through different LRR sensors are related is unclear.

In *Arabidopsis*, PRR LRR-RPs with molecularly defined ligands comprise receptors for bacterial, fungal or oomycete-derived necrosis and ethylene-inducing peptide 1-like proteins (NLPs) (RLP23)^4^; for proteobacterial translation initiation factor IF1 (RLP32)^18^ and for fungal polygalacturonases (PGs) (RLP42)^19,20^. In tomato (*Solanum lycopersicum*), fungal xylanases are sensed by the LRR-RP EIX2^21^, and in *Nicotiana benthamiana* bacterial cold shock protein is detected by CSPR^22^. Distribution of these immune receptors within the plant kingdom is often not only restricted to individual genera or even species, but some appear to exhibit significant sequence polymorphism within species. One example is RLP42, which exhibits remarkable plant accession specificity as active alleles were found in less than a third of 50 *Arabidopsis* accessions tested^20^, highly reminiscent of the pattern seen for many NLR alleles^16,17^. Tomato Ve1, Cf-2, Cf-4, Cf-4E and Cf-9 receptors are polymorphic plasma membrane-resident LRR-RPs that mediate plant cultivar-specific immunity through recognition of race-specific effectors produced by *Verticillium dahliae* (*Vd*Ave1) and *Cladosporium fulvum* (*Cf*Avr2, *Cf*Avr4, *Cf*Avr4E, *Cf*Avr9) fungal pathovars^23–28^. Thus, LRR-RP immune receptor family members are sensors for widespread microbial patterns and for microbial pathovar-specific effectors. LRR-RPs differ in their abilities to confer full immunity to infection. While RLP23, RLP32 and CSPR confer low-level (basal), PTI-type immunity to microbial infection, Ve1 and all identified *Cf*s provide ETI-like, complete immunity to fungal races producing the corresponding effectors^4,18,22,24,26,29^.

As plant LRR-RPs exhibit characteristics of sensors for both microbial surface patterns and polymorphic pathogen effectors, we investigated potential mechanistic overlaps in the architecture of immune signaling pathways governing PTI and ETI activation. Here we report that *Arabidopsis* cell surface LRR-RP immune receptors share with cytoplasmic NLR receptors an essential requirement for the EDS1-PAD4-ADR1 node in activating pattern- and effector-triggered immunity.

## Results

### LRR-RP-dependent immunity requires the RLCK-VII PBL31

We have reported opposing roles of the *Arabidopsis* RLCK-VII BIK1 in PTI conferred by LRR-RK or LRR-RP PRRs^7^. BIK1 negatively regulates LRR-RP-mediated PTI, but has a positive role in LRR-RK-mediated PTI^7,30,31^. BIK1 is released from the ligand-activated LRR-RK BAK1 complex and phosphorylates downstream targets to promote PTI^32–36^. We wanted to learn whether there are not only RLCK clade VII members with negative roles in LRR-RP-dependent PTI, but also RLCK-VII clade members with positive regulatory roles. We therefore screened a RLCK-VII T-DNA mutant library^37^ for ethylene production, a robust LRR-RP signaling output, elicited by three patterns recognized by LRR-RP PRRs (fungal pg13, recognized by RLP42^20^; microbial nlp20, recognized by RLP23^4^; bacterial eMax, recognized by RLP1^38^) (Fig. S1).

A *pbl31* mutant (SAIL_273_CO1) was consistently defective in its response to each LRR-RP elicitor (Fig. 1a, Fig. S1-S2), but not to the LRR-RK FLAGELLIN SENSING 2 (FLS2) elicitor, flg22, or the LRR-RK ELONGATION FACTOR RECEPTOR (EFR) elicitor elf18 (Fig. 1a, Fig. S2). PBL31 (PBS1-LIKE31) belongs to RLCK-VII subfamily 7, together with PBL30/CAST AWAY, which interacts with SOBIR1 during floral abscission^39^, and PBL32.^37^ In response to nlp20, *pbl30*, but not *pbl32* mutants, also produced slightly less ethylene than Col-0 wild-type (Fig. 1a). Ethylene production in a *pbl30 pbl31 pbl32* triple mutant or a *pbl30 pbl31* double mutant was reduced to a larger extent than in any single mutant. Collectively, these data point to a major role of PBL31 in LRR-RP-mediated signaling with minor contributions by PBL30 (Fig. 1a, Fig. S2). Ethylene production was fully complemented by overexpression of PBL31 in the triple mutant background (Fig. S3). *ACS2* and *ACS6* encode rate-limiting aminocyclopropane-1-carboxylic acid synthases in ethylene biosynthesis^40^, and the *acs2 acs6* double mutant fails to produce any ethylene upon stimulation with flg22 or nlp20 (Fig. S4a). Importantly, nlp20-induced expression of *ACS2* and *ACS6* was abolished in *pbl30 pbl31 pbl32* plants (Fig. S4b). A PBL31^K201A^ mutant, which carries a mutation in the putative protein kinase ATP binding pocket, did not restore ethylene production in *pbl30 pbl31 pbl32* (Fig. S3), suggesting that PBL31 kinase activity is required for LRR-RP-mediated PTI. Composite data from ten independent experiments revealed that nlp20-induced ethylene production was strongly reduced in *pbl30 pbl31 pbl32* plants compared to a slight, but statistically significant, reduction in flg22-induced ethylene production (Fig. 1a).

**Fig. 1.**
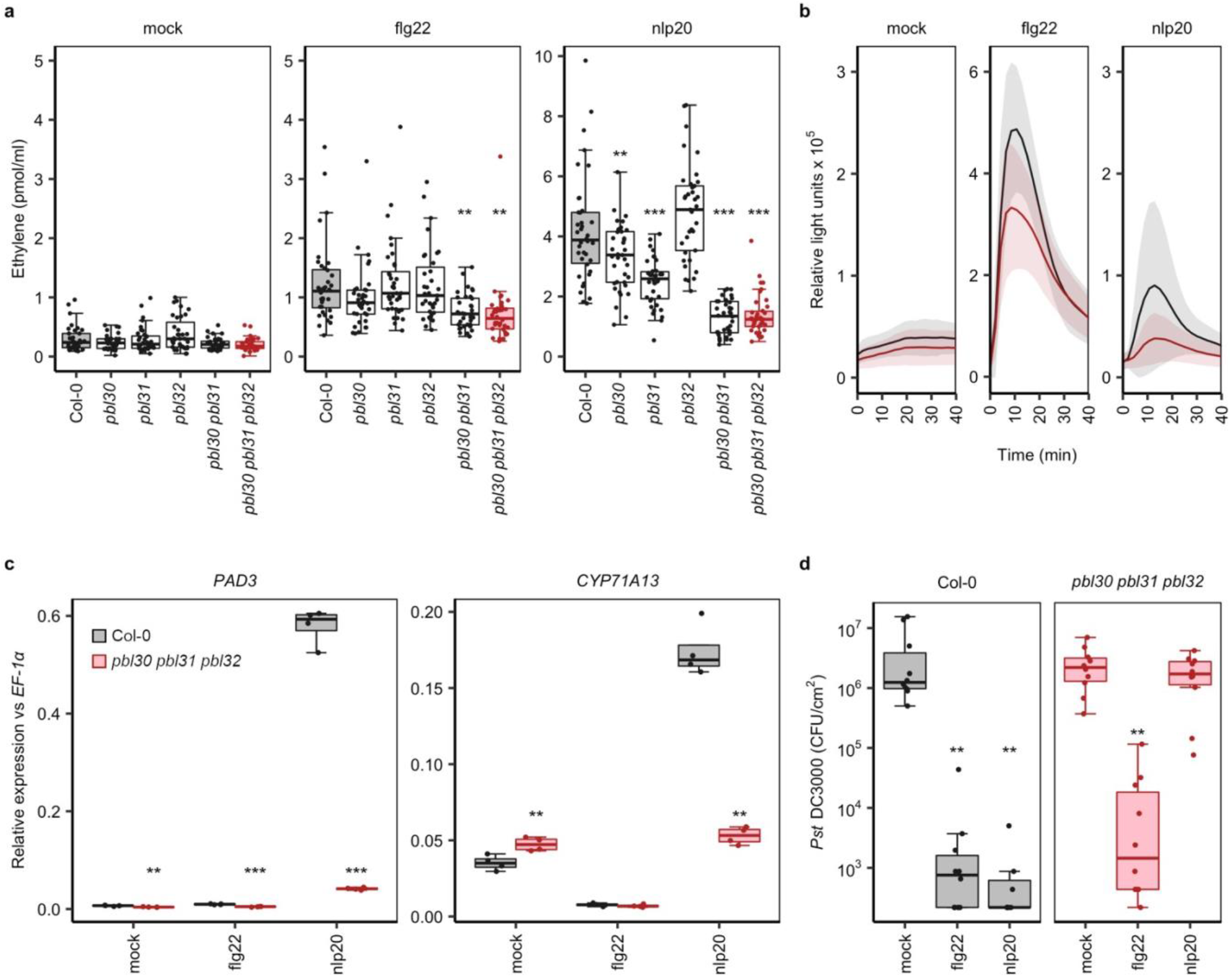
The role of RLCK-VII-7 family members in LRR-RP-mediated immunity. **a**, Ethylene accumulation in RLCK-VII-7 mutants treated with nlp20 or flg22. Leaf pieces of Col-0 and *pbl30 pbl31 pbl32* were treated with water or 500 nM of the indicated peptide. Ethylene accumulation was measured after 4 h. Composite data for 10 independent experiments (n=36) are shown. For all experiments, Col-0 is shown in grey and *pbl30 pbl31 pbl32* in red; all other mutants are shown as white boxes. **b**, Elicitor induced ROS production in Col-0 and *pbl30 pbl31 pbl32*. Leaf pieces of Col-0 and *pbl30 pbl31 pbl32* were treated with water (mock) or 500 nM of the indicated elicitor (n=16). The solid lines indicate the mean ROS response, and the shaded areas indicates standard deviation. **c**, Transcriptional profiling of camalexin biosynthesis genes by quantitative reverse transcription-PCR (qRT-PCR). Leaves of Col-0 or *pbl30 pbl31 pbl32* plants were infiltrated with water (mock) or 500 nM of the indicated elicitor and harvested after 6 h. Relative expression of the indicated genes was normalized to the levels of the *EF-1α*. Data are shown for one biological replicate with four technical replicates. The experiment was performed three times with similar results. **d**, Col-0 and *pbl30 pbl31 pbl32* leaves were infiltrated with 10 mM MgCl_2_ (mock), 1 μM nlp20 or 1 μM flg22. 24 h later, the plants were infiltrated with 10^4^ colony forming units (CFU) per mL of *Pst DC3000* suspended in 10 mM MgCl_2_. Bacterial colonization was determined after 3 days (n=10). For (**a**) and (**c**), asterisks indicate results of statistical tests for differences between the mutant and Col-0 response for the given elicitor (Dunnett’s test: ***, p<0.0001; **, p<0.01; *, p<0.05); for (**d**), asterisks indicate results of statistical test for differences between elicitor-primed and mock-treated samples (Steel’s test: **, p<0.01).

We examined whether the RLCK-VII-7 subfamily is required for other RLP23-mediated outputs. Nlp20-induced production of reactive oxygen species (ROS) in *pbl30 pbl31 pbl32* was virtually abolished, whereas flg22-induced ROS production was only slightly reduced (Fig. 1b, Fig. S5a). Also, nlp20-induced expression of the genes *PAD3* (*PHYTOALEXIN-DEFICIENT 3*) and *CYP71A13 (CYTOCHROME P71A13)*, encoding enzymes required for biosynthesis of the phytoalexin camalexin^41^, was impaired in *pbl30 pbl31 pbl32* leaves (Fig. 1c). These genes did not respond to flg22 in either wild-type or mutant plants (Fig. 1c)^7^.

PAMP-mediated priming of immunity to subsequent infection by a virulent pathogen is a characteristic of PTI^42,43^. We found that nlp20-induced priming was abolished whereas flg22-induced priming was not reduced in *pbl30 pbl31 pbl32* mutants (Fig. 1d, Fig. S6). In contrast, ETI conferred by the TIR-NLR receptor pair RRS1 RPS4 upon inoculation with the bacterial strain *Pseudomonas syringae* pv. *tomato* DC3000 AvrRPS4 was not diminished in *pbl30 pbl31 pbl32* (Fig. S7). We conclude that PBL31 is an essential positive regulator of LRR-RP SOBIR1-mediated PTI but not of LRR-RK-mediated PTI or TIR-NLR-mediated ETI.

### LRR-RP immunity is mediated by PAD4-EDS1 heterodimers

In *Arabidopsis*, lipase-like proteins EDS1, PAD4 and SAG101 constitute a key signaling node in ETI activation and basal immunity^12^. EDS1 forms exclusive heterodimers with either PAD4 or SAG101^44,45^. EDS1-SAG101 dimers are essential for TIR-NLR-dependent host cell death and immunity, whereas EDS1-PAD4 dimers principally trigger TIR-NLR and CC-NLR-dependent transcriptional defenses^12,13,46^. PAD4 and EDS1 are also involved in basal immunity, as *Arabidopsis eds1* and *pad4* mutants are hypersusceptible to virulent pathogens that lack strongly recognized effectors^47–49^. To test whether reduced basal immunity is due to impaired PTI, we tested PTI activation in a *pad4* mutant. In this genotype, we found substantially reduced levels of ethylene relative to those in Col-0 in response to LRR-RP ligands nlp20, IF1 and pg13 (Fig. 2a, Fig. S8). The low ethylene response mediated by LRR-RK FLS2 in Col-0 was slightly but statistically significantly reduced in *pad4*, as confirmed by composite data from 20 independent experiments (Fig. 2a). In contrast, LRR-RK EFR-mediated ethylene production was slightly, but not statistically significantly reduced (Fig. S8). To rule out possible regulatory effects of PAD4 on the expression of PRR immunity-associated genes, we assessed transcript levels of *SOBIR1*, *PBL31*, *RLP42* and *FLS2*. No substantial differences in the transcript levels of these genes were observed (Fig. S9a). Likewise, protein levels of BAK1 and FLS2 (the only *Arabidopsis* cell surface proteins involved in PTI for which specific and sensitive antisera are available) were unaltered in *pad4* relative to Col-0 (Fig. S9b), suggesting that expression and stability of the immunity-associated protein machinery is not affected by the lack of PAD4.

**Fig. 2.**
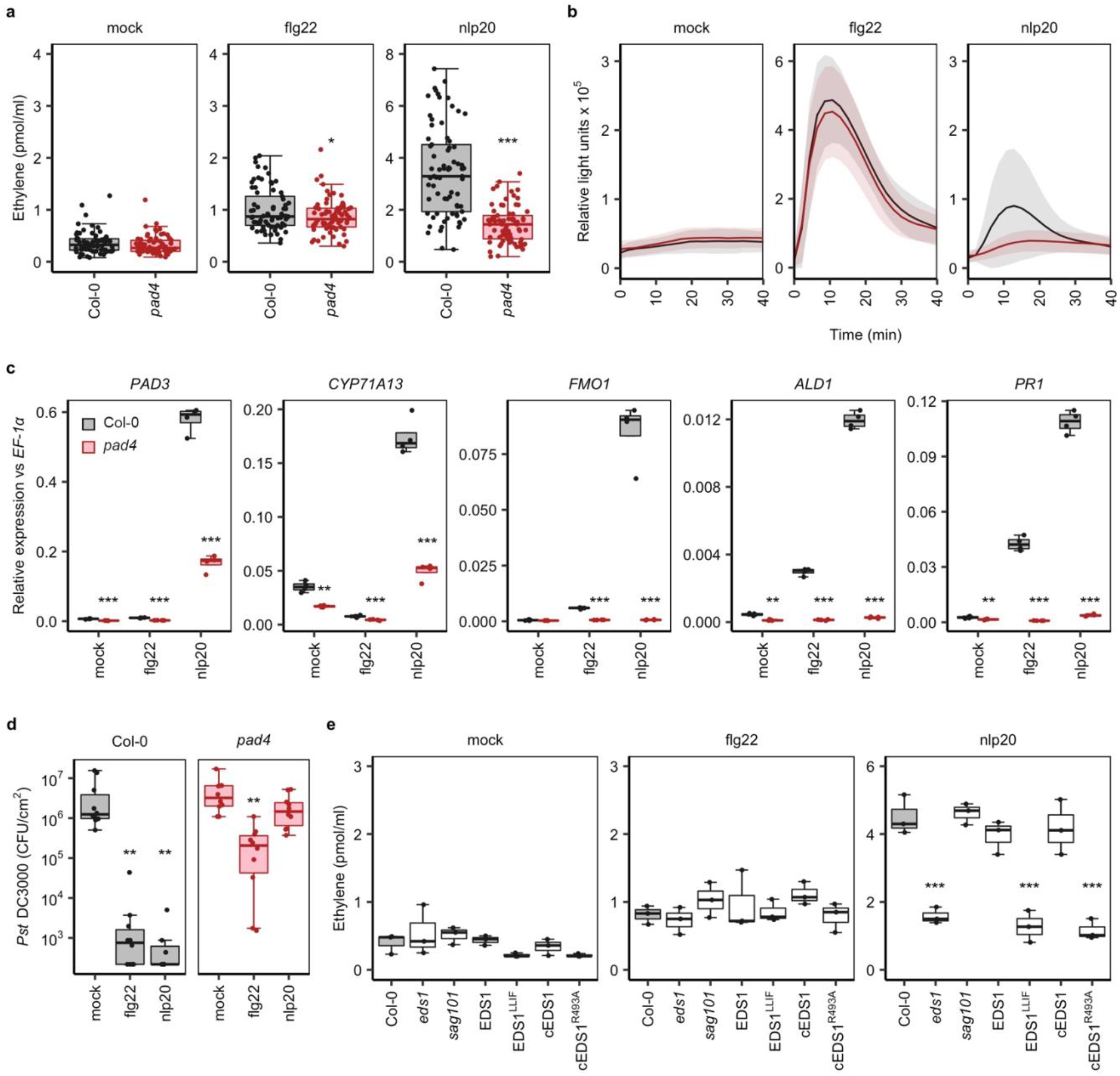
EDS1 and PAD4 are positive regulators of LRR-RP signaling. For all experiments, Col-0 is shown in grey and *pad4* in red. **a**, Nlp20-induced ethylene production is impaired in a *pad4* mutant. Leaf pieces of Col-0 and *pad4* were treated with water (mock) or 500 nM of the indicated peptide, and ethylene accumulation was measured after 4 h. Composite data for 20 independent experiments (n=72) are shown. **b**, Elicitor induced ROS production in Col-0 and *pad4*. Leaf pieces of Col-0 and *pad4* were treated with water (mock) or 500 nM of the indicated elicitor (n=16). The solid lines indicate the mean ROS response, and the shaded areas indicates standard deviation. **c**, Transcriptional profiling of *PAD3*, *CYP71A13*, *FMO1*, *ALD1* and *PR1* by qRT-PCR. Leaves of Col-0 or *pad4* plants were infiltrated with water (mock) or 500 nM of the indicated elicitor and harvested after 6 h. Relative expression of the indicated genes was normalized to the levels of the *EF-1α* transcript. Data represent one biological replicate with four technical replicates. The experiment was performed three times with similar results. **d**, Col-0 and *pad4* leaves were infiltrated with either 10 mM MgCl_2_ (mock), 1 μM nlp20 or 1 μM flg22. 24 h later, the plants were infiltrated with 10^4^ CFU/mL *Pst DC3000* suspended in 10 mM MgCl_2_. Bacterial colonization was monitored after 3 days (n=10). **e**, nlp20-induced ethylene response is dependent on an EDS1 PAD4 heterodimer signaling surface. The indicated lines were treated with water (mock), 500 nM nlp20 or 500 nM flg22. Ethylene accumulation was measured after 4 h (n=3). The experiment was performed 3 times with similar results. The *eds1* mutant is complemented with wild-type or mutant EDS1 or cEDS1 (from cDNA). For (**a**), (**c**) and (**e**), asterisks indicate results of statistical tests for differences between the mutant and Col-0 response for the given elicitor (Dunnett’s test vs Col-0: ***, p<0.0001; **, p<0.01; *, p<0.05); for (**d**), asterisks indicate results of statistical test for differences elicitor-primed and mock-treated samples (Steel’s test: **, p<0.01).

Our analysis of additional defense-related responses in *pad4* mutants revealed strongly reduced ROS production upon nlp20 but not flg22 treatment (Fig. 2b). Moreover, induction of *PAD3* and *CYP71A13* gene expression upon nlp20 treatment was reduced in *pad4* (Fig. 2c). These data suggest that PAD4 contributes to the activation of early and late PTI responses.

PAD4 is also a key regulator of systemic acquired resistance (SAR)^50^. SAR activation requires pipecolic acid and *N*-hydroxy-pipecolic acid that are produced by ALD1 (AGD2-LIKE DEFENSE RESPONSE PROTEIN 1) and FMO1 (FLAVIN-DEPENDENT MONOOXYGENASE)^51–55^. Pattern-induction of the *ALD1*, *FMO1* and the SA marker gene *PR1* (PATHOGENESIS-RELATED1) was reduced in *pad4* irrespective of the elicitor tested (Fig. 2c). The fungal phytotoxin thaxtomin A (TA) selectively activates PAD4-dependent immunity^56^. We found that TA pre-treatment of wild-type but not of a *pad4* mutant enhanced LRR-RP-but not LRR-RK-mediated ethylene production (Fig. S10), thus confirming a predominant involvement of PAD4 in LRR-RP signaling. Furthermore, nlp20 could no longer prime resistance to *Pst* DC3000 bacterial infection in the *pad4* mutant, whereas flg22-induced priming was only partially impaired (Fig. 2d, Fig. S6).

Most processes in which PAD4 is involved also require EDS1^44,45,49,57^. Consistent with this, we found that an *eds1* null mutant was deficient in LRR-RP-mediated responses, whereas a *sag101* null mutant responded similarly to the wild-type control (Fig. 2e, Fig. S8). To further elucidate the role of PAD4 with EDS1 in LRR-RP signaling, we tested whether interaction between the two proteins is necessary for RLP23 signaling. An *eds1* line complemented with an EDS1 variant (EDS1^LLIF^) that cannot dimerize with PAD4^44^ failed to restore the LRR-RP-mediated ethylene response (Fig. 2e). Likewise, mutation of a positively charged R493 residue (EDS1^R493A^) at the surface of a cavity formed by the EDS1 and PAD4 C-terminal domains disables ETI^46^ and reduced RLP23 signaling (Fig. 2e). Putative α/β-hydrolase catalytic residues in PAD4 and EDS1 N-terminal domains are dispensable for ETI and basal immunity^44,58^ and are also not required for the nlp20-induced ethylene response (Fig. S11). Thus, a stable PAD4-EDS1 heterodimer is required for LRR-RP SOBIR1-triggered immunity whereas the EDS1 and PAD4 putative catalytic residues are not. These observations are consistent with the established requirements for PAD4 and EDS1 in basal resistance and ETI. We conclude that an EDS1-PAD4 complex is essential not only for many aspects of NLR-mediated ETI, but also for LRR-RP-mediated PTI and in part for LRR-RK signaling. In contrast, the EDS1-SAG101 complex is exclusively used in ETI.

### LRR-RP signaling is mediated by ADR1 hNLRs

Helper NLRs are components of immune signaling networks downstream of sensor NLRs^11^. In *Arabidopsis*, EDS1 SAG101 dimers together with NRG1-family hNLRs form a signaling module that promotes host cell death in TIR-NLR ETI^10,13^. By contrast, EDS1-PAD4 heterodimers function together with ADR1-family hNLRs in TIR-NLR ETI, CC-NLR ETI and basal immunity^10,13^. We therefore tested contributions of these two hNLR families to LRR-RP triggered immunity. Pattern-triggered ethylene production was normal in an *NRG1* family double mutant (*nrg1 double*)^14^ (Fig. 3a, Fig. S12). In contrast, an *ADR1* family triple mutant (*adr1 triple*)^59^ did not produce ethylene when treated with nlp20, IF1 or pg13, while the response to flg22 or elf18 was only slightly attenuated (Fig. 3a, Fig. S12). A higher order *helperless* mutant^10^ with all *ADR1* and *NRG1* genes mutated behaved similarly to the *adr1 triple* mutant (Fig. 3a, Fig. S12). ROS production in *adr1 triple* plants was reduced upon nlp20 treatment, but not in response to flg22 (Fig. 3b). Many *Arabidopsis* sensor NLRs rely on RAR1 (REQUIRED FOR MLA (MILDEW LOCUS A) 12-MEDIATED RESISTANCE 1) and NDR1 (NON RACE-SPECIFIC DISEASE RESISTANCE 1) for proper accumulation and immune function, although this is not the case for ADR1-L2^60,61^. We found that RAR1 and NDR1 are also dispensable for pattern-induced ethylene production (Fig. S13).

**Fig. 3.**
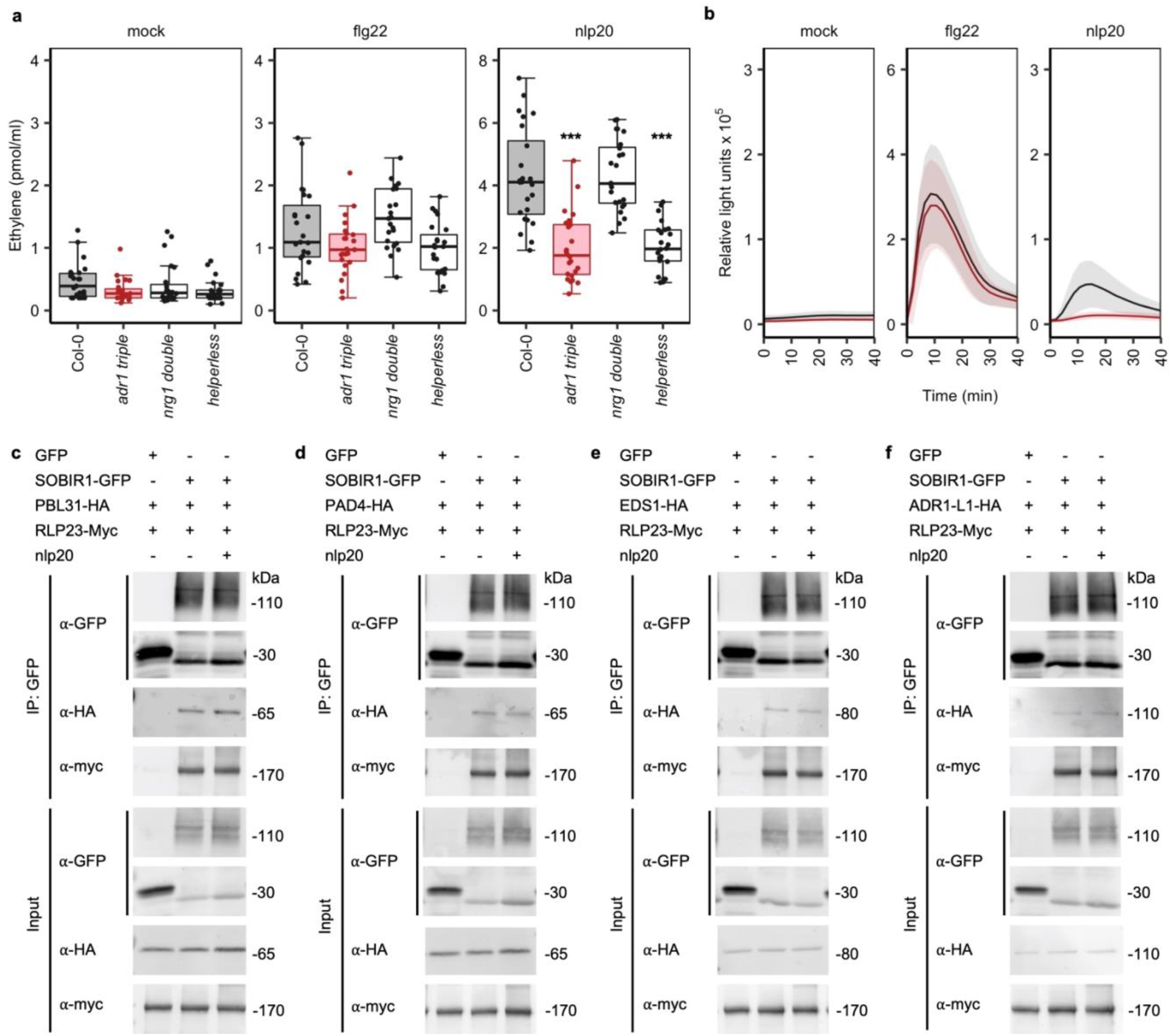
ADR1 helper NLRs are positive regulators of LRR-RP signaling and associate with a potential SOBIR1-PBL31-EDS1-PAD4 signaling node. For **a** and **b**, Col-0 is shown in grey, and *adr1 triple* is in red. **a**, Ethylene accumulation is impaired in higher order *adr1* and *helperless* mutants. Leaf pieces of the indicated lines were treated with either water (mock), 500 nM nlp20 or 500 nM flg22. Ethylene accumulation was measured after 4 h. Data from 6 independent experiments are shown (n=23). Asterisks indicate results of statistical tests for differences between the mutant and Col-0 response for the given elicitor (Dunnett’s test: ***, p<0.0001; **, p<0.01). **b**, Elicitor-induced ROS production in Col-0 and the *adr1* triple mutant. Leaf pieces of the indicated lines were treated with water (mock) or 500 nM of the indicated elicitor (n=16). The solid lines indicate the mean ROS response, and the shaded areas indicates standard deviation. **c-f**, (**c**) PBL31, (**d**) PAD4, (**e**) EDS1 and (**f**) ADR1-L1 associate with SOBIR1 in an nlp20-independent manner. SOBIR1-GFP and RLP23-Myc were transiently co-expressed with PBL31-HA, PAD4-HA or ADR1-L1-HA in *Nicotiana benthamiana.* Leaves were infiltrated with water or 1 μM nlp20, harvested after 10 min and subjected to co-immunoprecipitation with GFP-trap beads. Precipitated protein complexes were analyzed by protein blotting using tag-specific antisera.

While NLR-dependent ETI is often associated with host cell death, PTI is not. Recognition of fungal polygalacturonases by RLP42 provides a rare exception^19^, as treatment of Col-0 plants with a polygalacturonase fragment (pg23), but not with the elicitor-inactive mutant pg23m1^20^, induced chlorosis and lesion formation (Fig. S14). This lesion formation was abrogated in *pbl30 pbl31 pbl32*, but not in *eds1 pad4 sag101* or *helperless* mutants (Fig. S14), suggesting that the processes underlying PTI-associated and NLR sensor-mediated cell death are not identical. Together, these data highlight a shared requirement for the EDS1-PAD4-ADR1 node in immune signaling initiated intracellularly by sensor NLRs and at the cell surface by LRR-RP receptors.

### SOBIR1 associates with PAD4, EDS1, ADR1 family members and PBL31 to form a signaling complex

Our study supports roles of PBL31 and the EDS1-PAD4-ADR1 node in LRR-RP-mediated PTI. We used transient over-expression assays in *Nicotiana benthamiana* to test whether RLP23-SOBIR1 receptor complexes at the plasma membrane are arranged in spatial proximity with any of these newly discovered components. Expression of RLP23 and SOBIR1 confers nlp20 sensitivity to *N. benthamiana* plants^4^. When co-expressed, C-terminally epitope-tagged RLP23 and SOBIR1 precipitated PBL31 independently of the presence of nlp20 peptide, suggesting a ligand-independent stable interaction between PBL31 and the constitutive RLP23-SOBIR1 receptor complex (Fig. 3c). Likewise, SOBIR1 interacted in a ligand-independent manner with PAD4, EDS1 and the ADR1 family (ADR1, ADR1-L1 and ADR1-L2) (Fig. 3d-f, Fig. S15). Bimolecular fluorescence complementation assays further supported a close proximal arrangement of SOBIR1 at the plasma membrane with ADR1-L1 and ADR1-L2, but not ADR1 (Fig. S15b-c). Altogether, our findings suggest that plasma membrane-resident LRR-RP-SOBIR1 receptors form a supramolecular positive regulatory complex with RLCK-VII PBL31 and the EDS1-PAD4-ADR1 node to mediate PTI.

### Arabidopsis LRR-RP and NLR gene families exhibit similar levels of sequence polymorphisms

Having established that LRR-RPs and NLR immune receptors share similar signaling mechanisms, we wanted to learn whether the similarities extended also to evolutionary patterns. Both within species and within populations, NLR genes are highly diverse, with both signatures of rapid and balancing evolution^16,17,62^. The diversity in NLR repertoire is matched by pathogens being highly polymorphic for pathovar-specific effectors. In contrast, *Arabidopsis* LRR-RP-type immune receptors such as RLP1, RLP23, RLP30, RLP32 and RLP42 all recognize widespread microbial surface patterns^18,19,38,63,64^. Because these confer much lower levels of immunity to infection with virulent isolates, they should likely experience much weaker selection. To gain insight into LRR-RP diversity, reads from 80 *Arabidopsis* accessions from the first phase of the 1001 Genomes project^65,66^ were mapped to the TAIR10 assembly of the *Arabidopsis* Col-0 reference genome. Genes were categorized as being conserved, having complex patterns of variation or exhibiting presence/absence polymorphisms according to the distribution of large-scale polymorphisms across all accessions, as inferred from stringent read mappings. This profiling of within-species sequence diversity revealed that there is a similar fraction of variable genes within the NLR and LRR-RP gene families (Fig. 4a). By contrast, LRR-RK genes are much more conserved, comparable to variation in the genomic background (Fig. 4a). Intriguingly, LRR-RP genes encoding known PRRs were found in all three classes: RLP23, RLP30 and RLP32 showed a conserved pattern; RLP42 a complex pattern and RLP1 was characterized by presence/absence polymorphism (Supplementary Table 1). We concluded from this analysis that LRR-RP genes share with NLRs not only a genomic organization into gene clusters^67^ but also apparently similar evolutionary dynamics maintaining large sequence diversity (Fig. 4a), while LRR-RK-encoding genes are much more uniform.

**Fig. 4.**
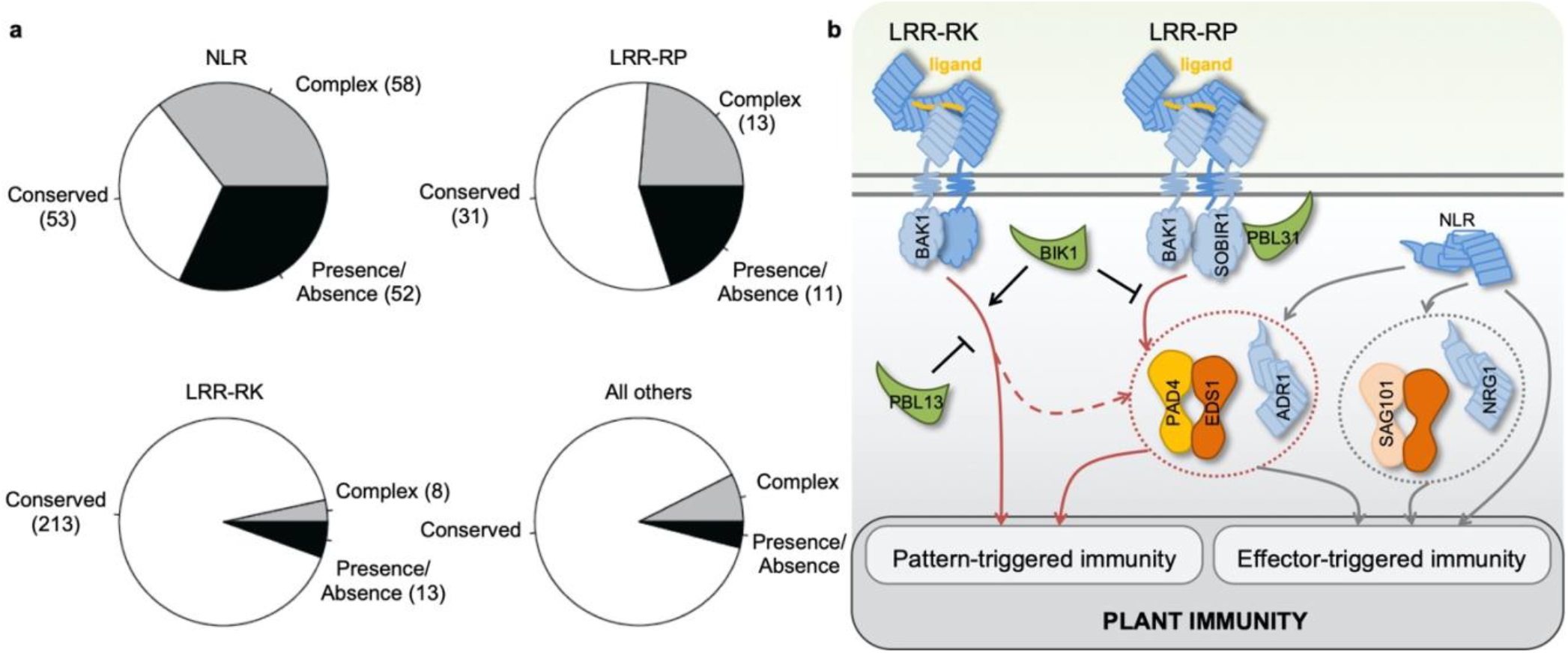
Conservation of NLR, LRR-RP and LRR-RK receptors and model of their convergence on the EDS1-PAD4-ADR1 signaling node. **a,** Fractions of NLR (163), LRR-RP (55) and LRR-RK (234) genes with conserved, presence/absence or complex patterns of variation. Categories were assigned based on the fraction of reference gene sequences covered with short reads from 80 *Arabidopsis* accessions. Numbers in parentheses indicate the number of genes in each category. **b**, Model depicting the EDS1-PAD4-ADR1 signaling node as a key mediator of both cell surface and intracellular immune signaling. Upon ligand perception, LRR-RK receptors form a complex with the co-receptor BAK1 to activate pattern-triggered immunity (PTI), which partially requires the EDS1-PAD4-ADR1 signaling node. By contrast, the LRR-RP-SOBIR1-BAK1 tripartite complex transduces the PTI signal mainly through the EDS1-PAD4-ADR1 node. Sensor NLRs activate effector-triggered immunity (ETI), which is dependent on the EDS1-PAD4-ADR1 and/or SAG101-EDS1-NRG1 nodes or independent of either signaling node. The RLCK-VII kinases (green) BIK1, PBL31 and PBL13 differentially regulate LRR-RP and LRR-RK signaling. PBL31 associates with SOBIR1 and plays a positive regulatory role in LRR-RP-mediated PTI, whereas BIK1 positively regulates LRR-RK-mediated PTI, but negatively regulates LRR-RP-mediated PTI. PBL13 has a negative role in LRR-RK-mediated PTI. Red arrows indicate PTI signaling and grey arrows indicate ETI signaling.

## Discussion

Plants employ two types of cell surface-resident, extracellular LRR domain immune receptors to sense proteinaceous ligands and trigger PTI: LRR-RKs and LRR-RPs^3,68,69^. Since both operate through ligand-induced recruitment of BAK1 to trigger immune signaling^2,3^ (Fig. 4b), an obvious hypothesis would be that they engage the same downstream pathways and that they are subject to similar evolutionary forces. However, the two PRR systems differ in their requirements for cytoplasmic RLCKs, and it has been proposed that the signaling pathways initiated by the two PRR types are mechanistically diverged^32,37^. BIK1 is a positive regulator of LRR-RK-mediated PTI, but has an opposite, negative regulatory role in LRR-RP-mediated PTI, in addition to its reported negative regulatory effects on aphid resistance and plant growth regulated by the hormone brassinolide^7,70,71^. PBL13 is a negative regulator of LRR-RK-mediated PTI^72^, but has no apparent role in LRR-RP-mediated immune activation (Fig. S1). We have now shown that RLCK clade VII-7 members PBL30 and PBL31 serve positive regulatory functions in LRR-RP-mediated defense activation (Fig. 1, Fig. S2). Thus, negative and positive regulation of PTI activation is brought about by specific RLCK family members, but in a remarkably PRR-type-dependent manner. It should be noted that RLCKs constitute one of several negative and positive regulatory mechanisms that control various facets of PTI activation in *Arabidopsis*^33,73–75^. We further report that PAD4 is essential for LRR-RP-dependent priming of immunity and activation of immunity-associated defense responses, but is only partially required for LRR-RK-dependent priming and defense activation (Fig. 2, Fig.8). This finding further substantiates the notion of different signal transduction cascades activated through different PRR systems^7^. Since both PRR systems, however, facilitate basal immune activation to microbial infection, it is assumed that their immune signaling networks display a rather high degree of functional redundancy or plasticity.

The EDS1-PAD4-ADR1 module has broad roles in ETI in both CC-NLR and TIR-NLR signaling pathways. We show that PTI activation in *Arabidopsis* mediated through the LRR-RP sub-class of cell surface immune receptors shares with intracellular NLR immune receptors the same essential molecular mechanistic requirement for the EDS1-PAD4-ADR1 node (Fig. 4b). The putative lipase activities of EDS1 and PAD4 are dispensable for immune activation, whereas the same EDS1-PAD4 heterodimer surface is essential for immune activation leading to either PTI or ETI (Fig. 2e). This finding has several implications: (i) Common use of an EDS1-PAD4-ADR1 node may provide an explanation for substantial overlaps in transcriptional and posttranscriptional defense output patterns observed upon activation of ETI or PTI^76^. (ii) Recently reported mutual potentiation of pattern and effector-triggered immune pathways suggests convergence of immune signaling pathways mediated through cell surface (PTI) and intracellular (ETI) immune receptors^77,78^. The EDS1-PAD4-ADR1 node at the plasma membrane is a candidate for linking cell surface and intracellular immune receptor signaling and might provide a molecular module through which mutual amplification of PTI and ETI could be brought about^76^. (iii) EDS1 and PAD4 have previously been implicated in basal host plant resistance to infection by host-adapted microbial pathovars^47–49^. Our findings indicate a role of these proteins in PTI, suggesting that pattern-induced defenses are a major component of basal immunity. (iv) Differential use of EDS1 and PAD4 may provide a molecular basis for maintaining signaling specificity in plant resistance to aphid and microbial infection. Unlike aphid resistance, in which PAD4 can act alone^58^, LRR-RP signaling requires *EDS1-PAD4* heterodimer formation. (v) The shared requirement of EDS1, PAD4 and ADR1 for PTI and ETI may also have evolutionary implications, suggesting that this node has primarily evolved to function in more ancient pattern-induced immunity.

PBL31, PAD4 and ADR1s have all been shown to interact with the LRR-RP co-receptor SOBIR1 in co-immunoprecipitation assays, thereby likely forming constitutive supramolecular immune signaling complexes at the inner side of the plant plasma membrane (Fig. 3c-f, Fig. S15). Like *Arabidopsis*, solanaceous plants may employ analogous modules shared between PTI and ETI signaling. Tomato EDS1 and solanaceous plant-specific hNLRs NLR REQUIRED FOR CELL DEATH 1 (NRC1) have been implicated in Ve1- and C*f*-4-dependent ETI, while Pto-dependent ETI and programmed cell death in *N. benthamiana* require NRC2a/b and NRC3^79–81^. NRC4 requirement for LRR-RP EIX2, but not for LRR-RK FLS2-mediated PTI activation in tomato has been reported^82^, and very recently, a gain-of-function mutation in NRC4a with increased basal resistance has been described^83^.

Cell surface LRR-RPs and cytoplasmic NLRs emerge as two highly polymorphic classes of immune sensors sharing an essential requirement for PAD4 in activation of inducible plant immunity (Fig. 4a). In contrast to NLRs, plant LRR-RP superfamily members comprise (i) conserved sensors for widely distributed microbial surface signatures, such as RLP23, CSPR or EIX2 ^4,21,22^, (ii) but also accession-specific, polymorphic sensors for widespread patterns, such as RLP42^19^, as well as (iii) sequence-divergent sensors for microbial pathovar-specific effectors, such as C*f* proteins^23–25,27^. LRR-RP receptors serve clearly distinguishable roles in host plant immunity as receptors conferring basal resistance to non-adapted pathogens (RLP23, CSPR, EIX2) and as receptors mediating full resistance to host-adapted microbial pathovars (C*f*-4, Ve1). Thus, members of the LRR-RP superfamily qualify not only as PRRs mediating PTI, but also as immune receptors mediating ETI. The functional dissection of LRR-RP-type immune receptors and their immunogenic ligands erodes the strict distinction between the two types of plant immunity^84^ and supports the view of plant immunity as a generic surveillance system for patterns of danger that are perceived by sets of surface-resident and intracellular immune receptors^85–87^. This concept is reinforced by the EDS1-PAD4-ADR1 node as an essential, shared element for immune activation through two classes of highly polymorphic immune sensors, LRR-RP cell surface receptors and cytoplasmic NLR receptors in *Arabidopsis*.

## Materials and Methods

### Plant material

Plant lines and mutants used in this study were all in the *Arabidopsis thaliana* accession Columbia-0 (Col-0) background and are listed in Supporting Information Table S2. Plants were grown in soil in climate chambers under short day conditions (8 h:16 h, light:dark, 150 μmol cm^−2^s^−1^ white fluorescent light, 40-60 % humidity, 22°C). *Nicotiana benthamiana* wildtype plants were grown in soil in either a greenhouse or climate chambers under 12 h:12 h light:dark cycle at 60-70 % humidity and 24-26°C.

### Elicitors used in this study

Flg22, elf18, nlp20, pg13, pg23 and pg23m1 peptides were synthesized according to the published sequences^20,43,88,89^ by Genscript Inc. (Piscataway, New Jersey, US), prepared as 10 mM stock solutions in DMSO and diluted in ddH2O prior to use. Full length IF1 from *E. coli* was synthesized by Genscript, Inc. as a biotinylated fusion protein and resuspended in ddH2O as a 1 mM stock solution^18^. The RLP1 elicitor eMax was originally identified in *Xanthomonas* (Jehle, et al)^38,90^. We found that eMax is also present in other proteobacteria including *Lysobacter*. Here we used eMax partially purified from a *Lysobacter* strain Root690^91^. Lysobacter was grown in SOB media overnight at 28°C with shaking at 200 rpm and harvested by centrifugation. The pellet was resuspended in 50 mM MES, pH 5.7, 50 mM NaCl, and cells were lysed by sonication, after which the supernatant was fractionated using a HiTrapQ FF (GE Healthcare, Uppsala, Sweden) anion exchange column. An eMax-containing fraction with high ethylene-inducing activity on *fls2 efr* leaves, but no activity on *rlp1* leaves was used for the RLCK-VII mutant screen as shown in Fig. S1.

### Measurement of reactive oxygen species (ROS) production

ROS assays were performed as described^88,92^. Leaves of 5-week old *Arabidopsis* plants were cut into pieces of equal size and floated on H2O overnight. One leaf piece per well was transferred to a 96-well plate containing 20 μM L-012 (Wako Pure Chemical Industries Ltd, Osaka, Japan) and 2 μg ml^−1^ peroxidase. Luminescence was measured over 1 h following elicitation or mock treatment using a Mithras LB 940 luminometer (Berthold Technologies, Bad Wildbad, Germany).

### Measurement of ethylene production

Leaves of 6-week old *Arabidopsis* plants were cut into pieces (~0.5 cm x 0.5 cm) and floated on H2O overnight. Three leaf pieces were incubated in a sealed 6.5 ml glass tube with 0.4 ml of 50 mM MES buffer, pH 5.7 and the indicated elicitor. Ethylene accumulation was measured after 4 h by gas chromatographic analysis (GC-14A; Shimadzu, Duisburg, Germany) of 1 ml of the air drawn from the closed tube with a syringe.

### Pathogenicity and cell death assays

*P. syringae* pv. *tomato* DC3000 (*Pst* DC3000) inoculations were performed as described^43^. For priming assays, leaves of 4- to 6-week old *Arabidopsis* plants were infiltrated with 1 μM nlp20, 1 μM flg22 or mock-treated 24 h prior to bacterial infection. Leaves were infiltrated with *Pst* DC3000 or *Pst* DC3000 AvrRPS4 at a density of 10^4^ cells/mL and bacterial growth was quantified after 3 d. For PTI-mediated cell death assays (Fig. S14), leaves of 5-6-week-old *Arabidopsis* plants were infiltrated with 10 μM pg23 or inactive pg23m1, and chlorosis was observed after 7 days.

### Quantitative reverse transcription PCR

Leaves from 6-week-old *Arabidopsis* plants were infiltrated with water (mock) or the indicated elicitors. Total RNA was isolated from leaves harvested at the indicated time point using the NucleoSpin^®^ RNA Plus kit (Macherey-Nagel, Düren, Germany). cDNA was synthesized from 2 μg of total RNA using the RevertAid First Strand cDNA Synthesis Kit (Thermo Scientific, Waltham, MA, USA). Quantitative PCR reactions and measurements were performed with a CFX384 Real-Time PCR detection system or an iQ5 Multi-color real-time PCR detection system (Bio-Rad, Hercules, CA, USA) using the SYBR Green Fluorescein Mix (Thermo Scientific) and the primers listed in Table S3. Transcript levels of target genes were normalized to the transcript levels of the housekeeping gene *EF*-*1α*.

### Immunoprecipitation

Leaves of *N. benthamiana* were transiently transformed with the indicated constructs and harvested after 2-3 days. For the elicitor-treated samples, leaves were infiltrated with water or 1 μM nlp20 10 min prior to harvesting. Immunoprecipitations were performed with 200-250 mg of tissue. Tagged proteins were immunoprecipitated for 1 h at 4°C using GFP-Trap beads (ChromoTek, Planegg, Bavaria, Germany) following the method of Chinchilla et al^5^ or for Fig. S15a, El Kasmi et al^93^. Protein blotting was performed using antibodies against GFP (Torrey Pines Biolabs, Secaucus, New Jersey, US), HA (Sigma, St. Louis, Missouri, US) or Myc (Sigma). For Fig. S15a, protein blotting was performed using antibodies against GFP (Roche, Basel, Switzerland) and HA (Roche).

### Ratiometric bimolecular fluorescence complementation (rBiFC)

The coding sequences of SOBIR1, ADR1, ADR1-L1 and ADR1-L2 were cloned into the 2in1 BiFC CC gateway-compatible destination vector^94,95^. Destination vectors were transiently expressed in *N. benthamiana* and complementation of yellow-fluorescence protein was analyzed at 24 hours after infection (hpi) with the confocal laser scanning microscope LSM880 (Zeiss, Oberkochen, Germany) using the 63x water-immersion objective. Settings were as follows: YFP was excited using a 514 nm laser, collecting emission between 516-556 nm; RFP was excited using a 561 nm laser with an emission spectrum of 597-634 nm. Images were processed with ZENblue software (Zeiss) for adjustment of brightness and contrast.

### Conservation analysis of LRR-RKs, LRR-RPs and NLRs in Arabidopsis

Reads from 80 *A. thaliana* accessions from the first phase study of the 1001 Genomes project^65,66^ were mapped to the reference genome of Col-0 using version 0.7.15-r1140 of the BWA-backtrack algorithm^96^ with parameters: *k*=1 in bwa aln command; *n*=10000; with maximal number of mismatches allowed 1. Paired end information was discarded. The TAIR10 assembly of the *A. thaliana* Col-0 genome was used for the reference genome (https://arabidopsis.org). The output mapped files were processed with samtools mpileup version 1.9^97^; parameters: *aa*; *d*=10000; *Q*=0. A total list of 163 NLR genes was used in the analysis, which was based on 159 NLR genes previously identified^67^ and four additional, manually curated genes (*AT1G63860*, *AT1G72920*, *AT1G72930* and *AT5G45230*). The coding sequence (CDS) portions of the genes were extracted, defined as the overlap of all the CDS models of the gene based on TAIR10 annotation. Fractions of CDS sequence with non-zero coverage were calculated for each gene-accession combination (hereafter “coverage fractions”). Genes were assigned into conserved, presence/absence and complex categories using a threshold-based approach. To define thresholds, *k* means algorithm was initiated with three centers at 0, 0.5 and 1 and applied to the coverage fractions, resulting in thresholds at 0.37 and 0.81. Coverage fractions were then discretized by applying these thresholds into ‘absent’, ‘intermediate’ and ‘present’ categories, from lowest to highest values. NLR genes were assigned as conserved if there were no accessions with ‘absent’ coverage and at least 95% of all accessions had high coverage. Genes with more than 5% of accessions having ‘intermediate’ coverage values were assigned as complex, and genes which were absent in at least one accession not classified as complex, were assigned as presence/absence. This procedure was also applied to LRR-RPs^98^ and receptor-like kinases (RKs) including LRR-RK-encoding genes^99^. The conserved category does not necessarily imply functional or structural conservation, but is used in the genomic sense to indicate sequence conservation, as measured by the presence of sub-sequences whose identities are within the applied thresholds.

### Statistical analysis

Data sets were analyzed using Microsoft Office Excel, R or JMP. Comparisons with the control were made using Dunnett's test. For priming assays (Fig. 1d, Fig. 2d and Fig. S6), the data showed a nonparametric distribution and were therefore analyzed using Steel’s test.

## Acknowledgments

This work was supported by Deutsche Forschungsgemeinschaft (DFG) grants Nu 70/15-1, ERA-CAPS-Grant SICOPID Nu 70/16-1 to T.N.; grant CRC-1101 to F.E.K., D.W. and T.N. and grant CRC-1403-414786233 to J.E.P. S.C.S. was supported by the Reinhard Frank Stiftung (Project ‘helperless plant’). F.L., J.E.P., D.K. and D.W. were supported by the Max-Planck-Society. We thank Sonja Harter for assisting in the generation of the ADR1 and ADR1-L1 BiFC entry constructs and Eunyoung Chae for annotation information of NLRs.

## Supplemental Data

**Table S1.** Classification of LRR-RPs according to genetic conservation in 80 *Arabidopsis* accessions (corresponds with Fig. 4A).

| **RLP** | **Gene ID** | **Classification** |
| --- | --- | --- |
| *RLP1* | AT1G07390 | Presence/Absence |
| *RLP2* | AT1G17240 | Presence/Absence |
| *RLP3* | AT1G17250 | Presence/Absence |
| *RLP4* | AT1G28340 | Conserved |
| *RLP5* | AT1G34290 | Complex |
| *RLP6* | AT1G45616 | Conserved |
| *RLP7* | AT1G47890 | Complex |
| *RLP8* | AT1G54470 | Conserved |
| *RLP9* | AT1G58190 | Conserved |
| *RLP10* | AT1G65380 | Conserved |
| *RLP11* | AT1G71390 | Presence/Absence |
| *RLP12* | AT1G71400 | Conserved |
| *RLP13* | AT1G74170 | Complex |
| *RLP14* | AT1G74180 | Conserved |
| *RLP15* | AT1G74190 | Conserved |
| *RLP16* | AT1G74200 | Presence/Absence |
| *RLP17* | AT1G80080 | Conserved |
| *RLP19* | AT2G15080 | Conserved |
| *RLP20* | AT2G25440 | Conserved |
| *RLP21* | AT2G25470 | Conserved |
| *RLP22* | AT2G32660 | Conserved |
| *RLP23* | AT2G32680 | Conserved |
| *RLP24* | AT2G33020 | Presence/Absence |
| *RLP25* | AT2G33030 | Presence/Absence |
| *RLP26* | AT2G33050 | Conserved |
| *RLP27* | AT2G33060 | Conserved |
| *RLP28* | AT2G33080 | Presence/Absence |
| *RLP29* | AT2G42800 | Conserved |
| *RLP30* | AT3G05360 | Conserved |
| *RLP31* | AT3G05370 | Conserved |
| *RLP32* | AT3G05650 | Conserved |
| *RLP33* | AT3G05660 | Conserved |
| *RLP34* | AT3G11010 | Complex |
| *RLP35* | AT3G11080 | Conserved |
| *RLP36* | AT3G23010 | Presence/Absence |
| *RLP37* | AT3G23110 | Complex |
| *RLP38* | AT3G23120 | Complex |
| *RLP39* | AT3G24900 | Complex |
| *RLP40* | AT3G24982 | Complex |
| *RLP41* | AT3G25010 | Complex |
| *RLP42* | AT3G25020 | Complex |
| *RLP43* | AT3G28890 | Complex |
| *RLP44* | AT3G49750 | Conserved |
| *RLP45* | AT3G53240 | Conserved |
| *RLP46* | AT4G04220 | Conserved |
| *RLP47* | AT4G13810 | Complex |
| *RLP48* | AT4G13880 | Presence/Absence |
| *RLP50* | AT4G13920 | Presence/Absence |
| *RLP51* | AT4G18760 | Conserved |
| *RLP52* | AT5G25910 | Conserved |
| *RLP53* | AT5G27060 | Conserved |
| *RLP54* | AT5G40170 | Conserved |
| *RLP55* | AT5G45770 | Conserved |
| *RLP56* | AT5G49290 | Complex |
| *RLP57* | AT5G65830 | Conserved |

**Table S2.** *Arabidopsis* lines used in this study.

| Line | Locus | Description | Reference |
| --- | --- | --- | --- |
| *acs2* | At1g01480 | Insertion, *acs2-1* | Tsuchisaka et al. (2009) |
| *acs6* | At4g11280 | Insertion, *acs6-1* | Tsuchisaka et al. (2009) |
| *acs2 acs6* | At1g01480, At4g11280 | *acs2-1/acs6-1* double mutant | Tsuchisaka et al. (2009) |
| *adr1 triple* | At1g33560, At4g33300, At5g04720 | Triple mutant of *adr1-1* (SAIL_842_B05), *adr1-L1-1* (SAIL_302_C06) and *adr1-L2-4* (SALK_126422) | Bonardi et al. (2011) |
| *eds1* | At3g48090 | Polymorphism 1009135505, *eds1-2*, introgressed into Col-0 | Bartsch et al. (2006) |
| *eds1 cEDS1* | At3g48090 | Col-0 *eds1-2* complemented with *cEDS1* (cloned from cDNA) | Bhandari et al. (2019) |
| *eds1 cEDS1<sup>R493A</sup>* | At3g48090 | Col-0 *eds1-2* complemented with *cEDS1<sup>R493A</sup>*, harboring a mutation in a conserved EP-domain surface in by EDS1/PAD4 dimers | Bhandari et al. (2019) |
| *eds1 EDS1* | At3g48090 | Col-0 *eds1-2* complemented with EDS1 (cloned from gDNA) | Wagner et al. (2013) |
| *eds1 EDS<sup>LLIF</sup>* | At3g48090 | Col-0 *eds1-2* complemented with EDS1<sup>LLIF</sup>, a mutant impaired in PAD4 and SAG101 dimerization | Wagner et al. (2013) |
| *eds1 pad4* | At3g52430, At3g48090 | Col-0 *eds1-2, pad4-1* double mutant | Wagner et al. (2013) |
| *eds1 pad4 EDS1 PAD4* | At3g52430, At3g48090 | Col-0 *eds1-2pad4-1* mutant complemented with wild-type EDS1 and PAD4 | Wagner et al. (2013) |
| *eds1 pad4 EDS1<sup>SDH</sup> PAD4<sup>S</sup>* | At3g52430, At3g48090 | Col-0 *eds1-2pad4-1* mutant complemented with EDS1 and PAD4 with mutations in the predicted catalytic residues | Wagner et al. (2013) |
| *eds1 pad4 sag101* | At3g52430, At3g48090, At5g14930 | Col-0 *eds1-2, pad4-1, sag101-2* triple mutant | Wagner et al. (2013) |
| *helperless* | At1g33560, At4g33300, At5g04720, At5g66900, At5g66910, At5g66890 | Pentuple mutant of all full-length and functional *ADR1* and *NRG1* genes in Col-0; *NRG1.1* and *NRG1.2* CRISPR/Cas9 deletion mutant in the triple insertion mutant of *adr1-1 adr1-L1-1 adr1-L2-4* | Saile et al. (2020) |
| *ndr1* | At3g20600 | Polymorphism 1005991898, *ndr1-1* | Century et al. (1995) |
| *nrg1 double* | At5g66900, At5g66910 | *NRG1.1* and *NRG1.2* CRISPR/Cas9 deletion mutant | Castel et al. (2019) |
| *pad4* | At3g52430 | Polymorphism, 4770301, *pad4-1* | Jirage, et al. (1999) |
| *pbl30* | At4g35600 | Insertion, SAIL_296_A06, also known as *cst-2* | Burr et al. (2011) |
| *pbl30 pbl31* | At4g35600, At1g76360, | Insertions, SAIL_296_A06, SAIL_273_C01 | This study |
| *pbl30 pbl31 pbl32* | At4g35600, At1g76360, At2g17220 | Triple mutant of SAIL_296_A06, SAIL_273_C01 and SALK_113804 | Rao et al. (2018) |
| *pbl30 pbl31 pbl32 PBL31* | At4g35600, At1g76360, At2g17220 | *pbl30 pbl31 pbl32* complemented with PBL31 | This study |
| *pbl30 pbl31 pbl32 PBL31<sup>K201A</sup>* | At4g35600, At1g76360, At2g17220 | *pbl30 pbl31 pbl32* complemented with the kinase dead mutant PBL31<sup>K201A</sup> | This study |
| *pbl31* | At1g76360 | Insertion, SAIL_273_C01 | Rao et al. (2018) |
| *pbl32* | At2g17220 | Insertion, SALK_113804 | Rao et al. (2018) |
| *rar1* | At5g51700 | Polymorphism 6530624064, *rar1-21* | Tornero et al. (2002) |
| *sag101* | At5g14930 | Insertion, *sag101-2* | Feys et al. (2005) |
| *sobir1* | At2g31880 | Insertion, SALK_050715, *sobir1-12* | Gao et al. (2009) |

**Table S3.**
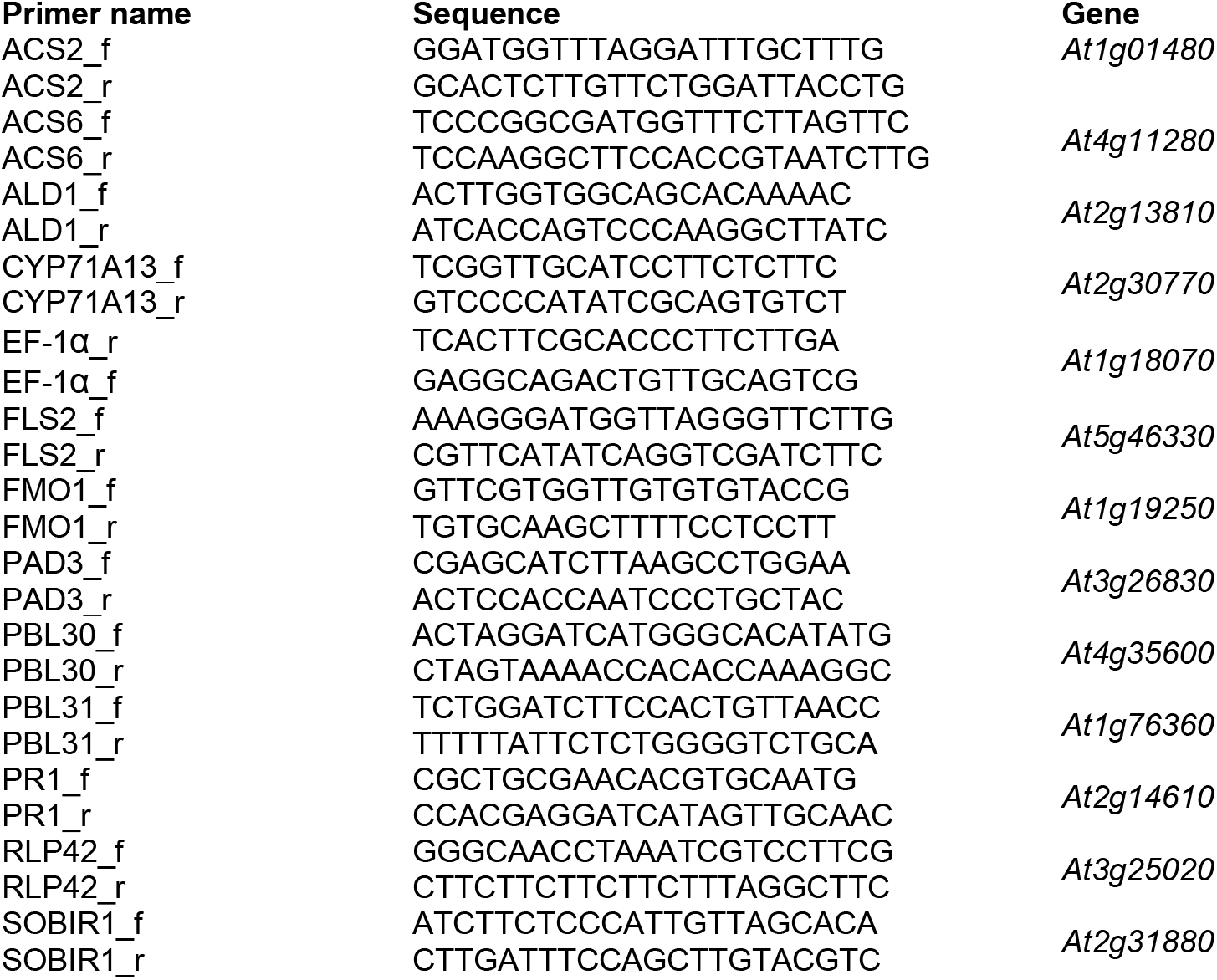
Primers used for quantitative RT-PCR.

**Fig. S1.**
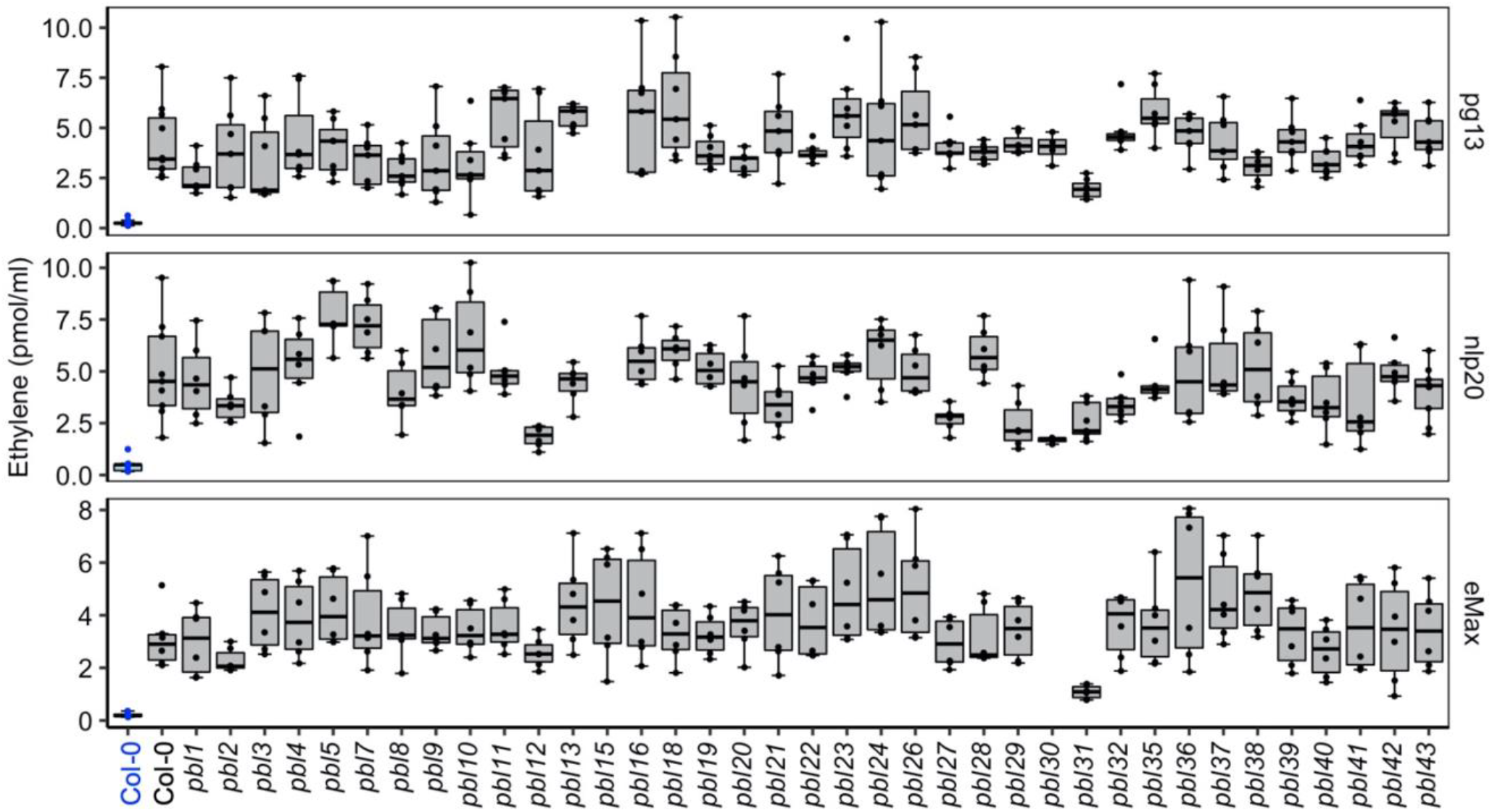
Response of *Arabidopsis* RLCK-VII mutant lines to LRR-RP elicitors. Leaf pieces of the indicated lines were treated 500 nM pg13, 500 nM nlp20 or a partially purified extract containing eMAX. Ethylene accumulation was measured after 4 h (n≥6). Mock treated Col-0 is shown in blue.

**Fig. S2.**
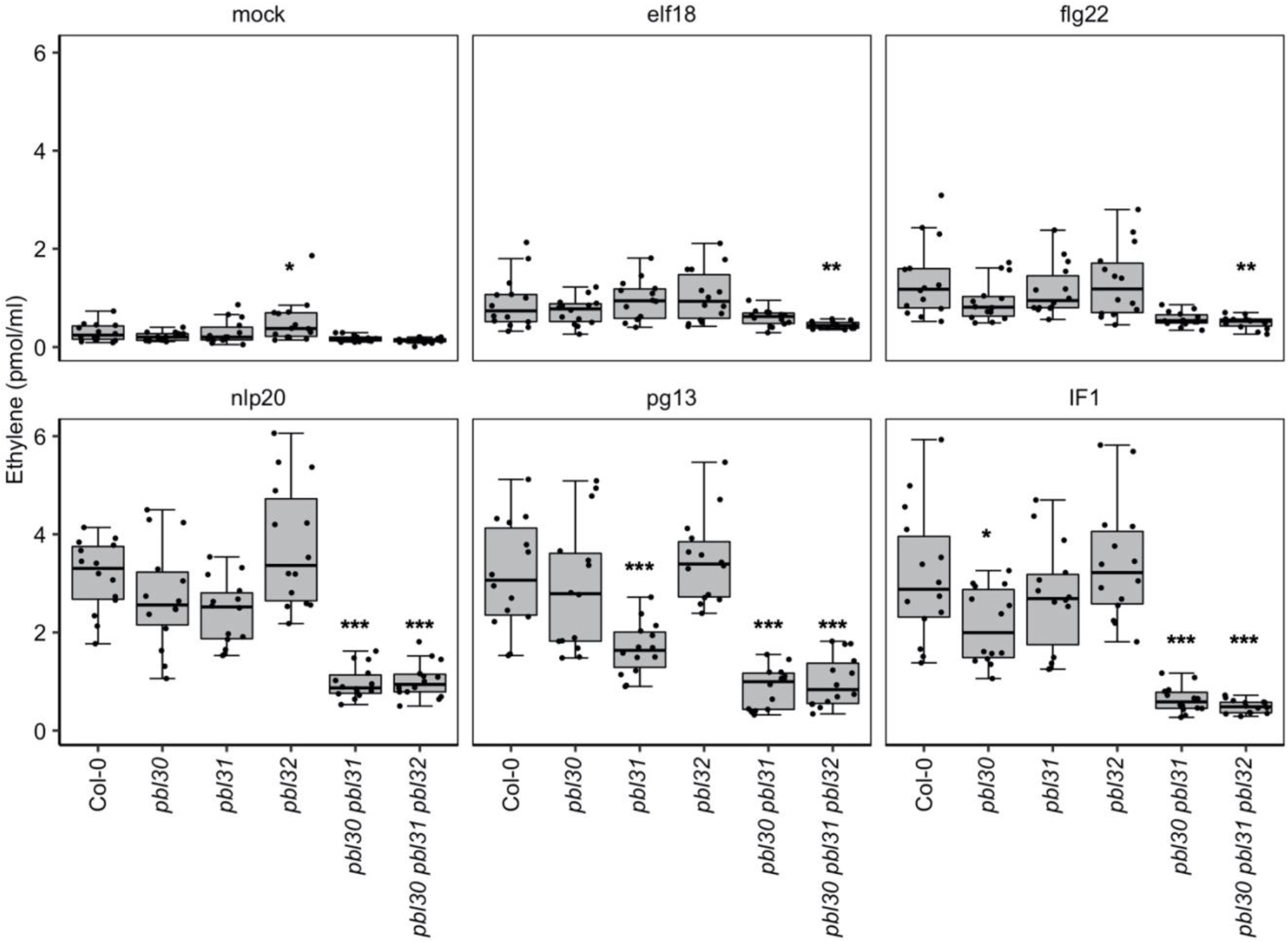
Ethylene accumulation in RLCK-VII-7 mutants treated with LRR-RP and LRR-RK elicitors. Leaf pieces of the indicated lines were treated with water (mock), 500 nM elf18, 500 nM flg22, 500 nM nlp20, 500 nM pg13 or 100 nM IF1. Ethylene accumulation was measured after 4 h. Data from three independent experiments (n=13 shown as box plots. Asterisks indicate results of statistical tests for differences between the mutant and Col-0 response for the given elicitor (Dunnett’s test vs Col-0: ***, p<0.0001; **, p<0.01; *, p<0.05).

**Fig. S3.**
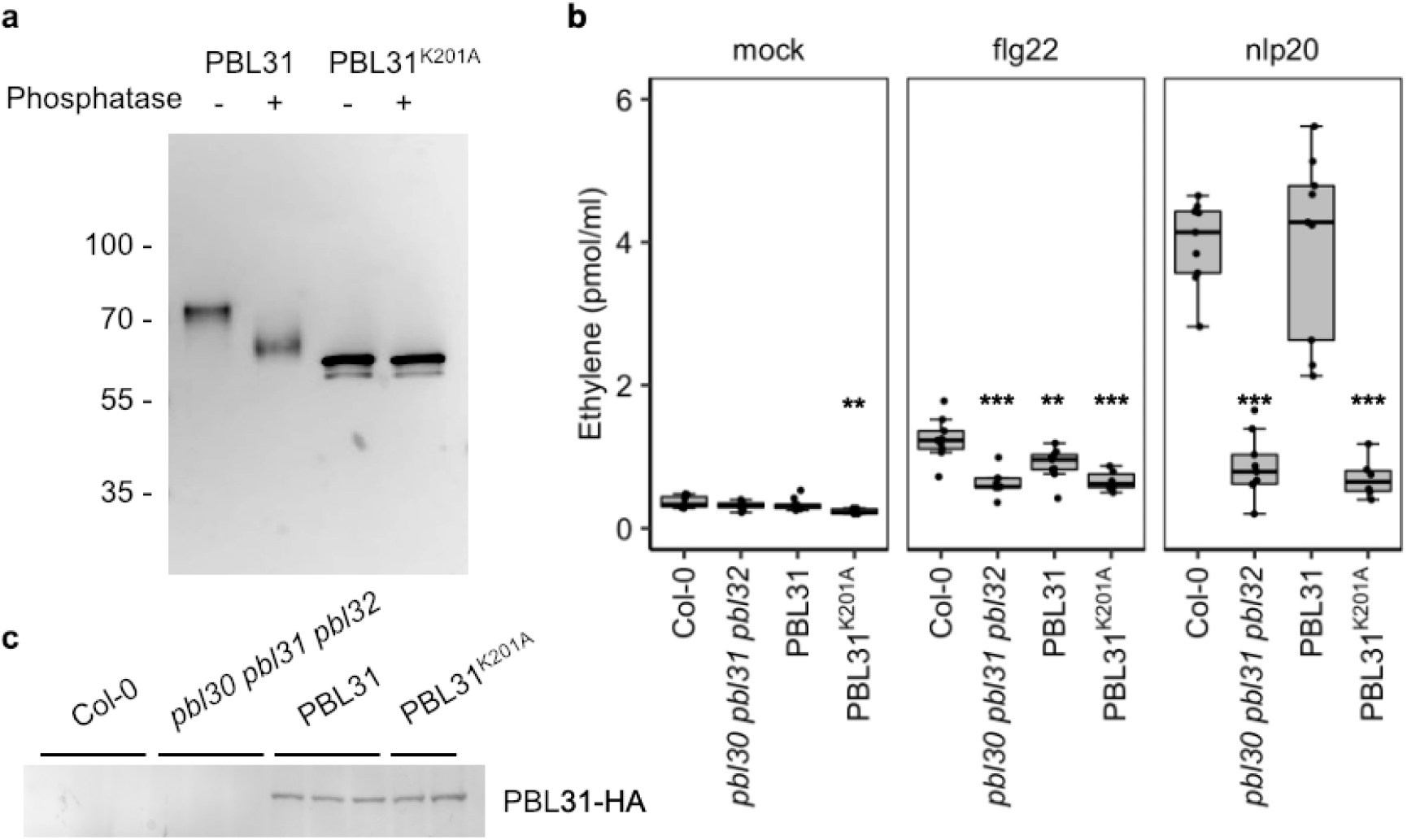
PBL31 activity in LRR-RP signaling requires kinase activity. **a**, PBL31 has autokinase activity that is abolished in the PBL31^K201A^ mutant. Recombinant PBL31 and PBL31^K201A^ were analyzed by anti-His protein blot. PBL31^K201A^ runs near the predicted position for the tagged protein (57.4 kDa). The wild-type version migrates more slowly, consistent with its being auto-phosphorylated. Phosphorylation of the wild-type PBL31 was confirmed by treatment with calf intestinal phosphatase, which increased the SDS-PAGE migration rate of PBL31 but not PBL31^K201A^. **b**, Ethylene accumulation in *pbl30 pbl31 pbl32* complemented with wild-type PBL31 or the kinase dead variant PBL31^K201A^. Leaf pieces of the indicated lines were treated with water (mock) or 500 nM of the indicated peptide. Ethylene accumulation was measured after 4 h (n≥6). Asterisks indicate results of statistical tests (Dunnett’s test vs Col-0: ***, p<0.0001; **, p<0.01). **c**, Anti-HA protein blot of plants used in panel (**b**).

**Fig. S4.**
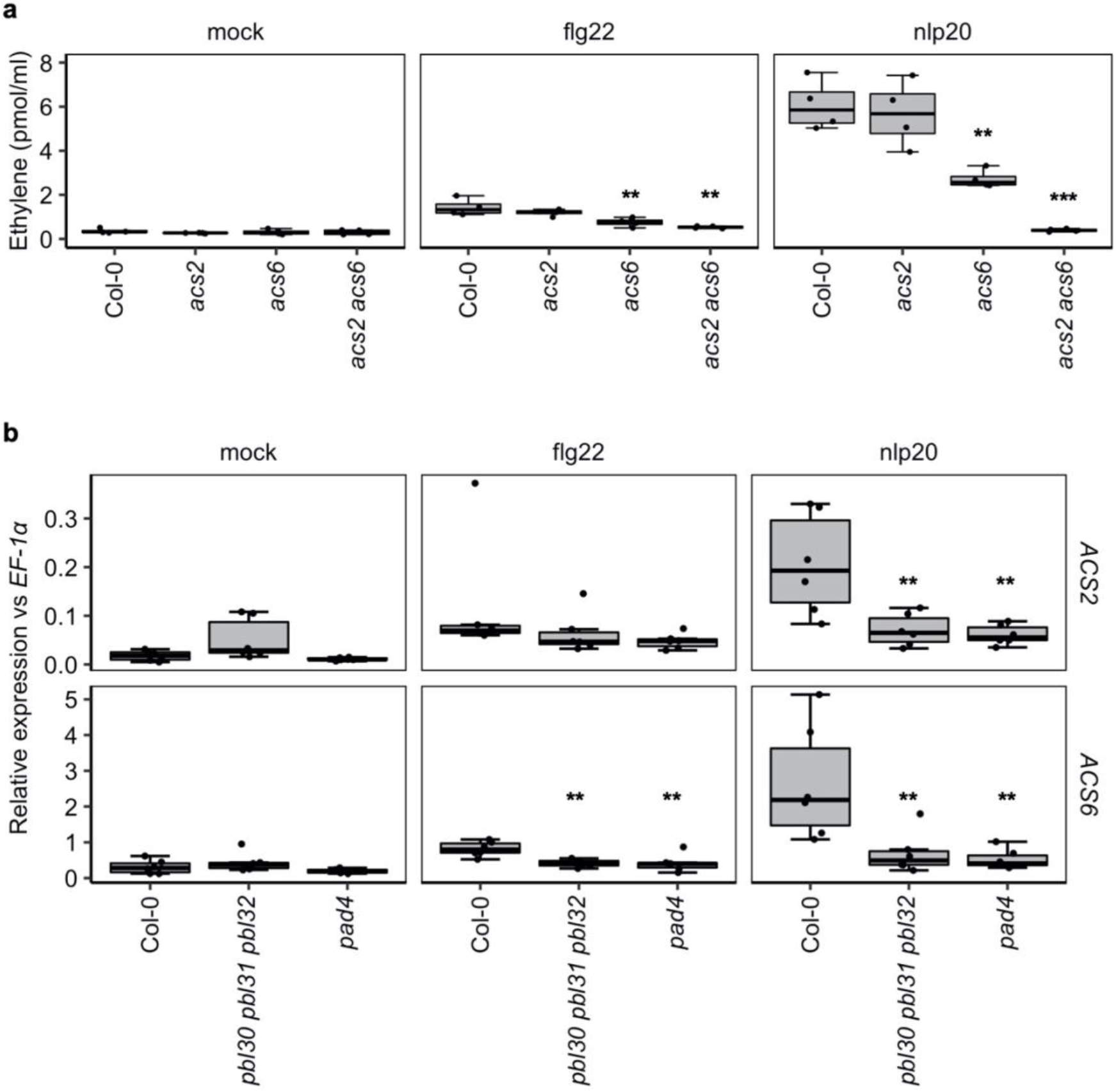
The role of the ACC synthase genes *ACS2* and *ACS6* in nlp20- and flg22-induced ethylene responses. **a**, *ACS2* and *ACS6* are critical for ethylene production in response to nlp20 and flg22. Col-0 and the indicated mutant lines were analyzed for their ability to accumulate ethylene in response to nlp20 and flg22. Leaf pieces of the indicated lines were treated with water (mock), 1 μM flg22 or 1 μM nlp20. Ethylene accumulation was measured after 4 h (n = 4). The experiment was repeated three times with similar results. **b**, Transcriptional profiling of *ACS2* and *ACS6* by quantitative reverse transcription-PCR (qRT-PCR). Leaves of Col-0, *pad4* or *pbl30 pbl31 pbl32* plants were infiltrated with water (mock) or 500 nM of the indicated elicitors and harvested after 1.5 h. Relative expression of the indicated genes is shown normalized to the *EF-1α* transcript. Data are from three biological replicates. Asterisks indicate results of statistical tests (Dunnett’s test vs Col-0: ***, p<0.0001; **, p<0.01).

**Fig. S5.**
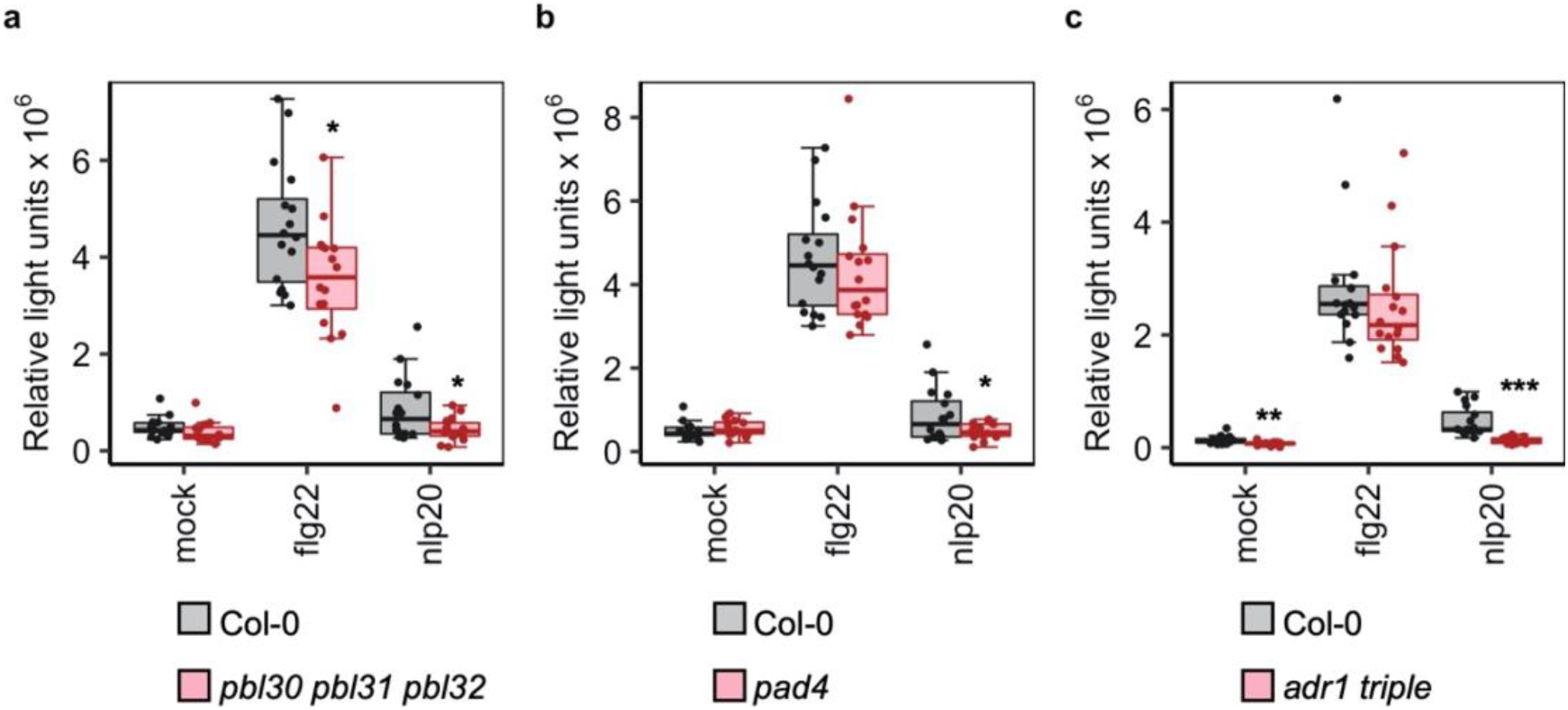
ROS production is impaired in *pbl30 pbl31 pbl32*, *pad4* and *adr1 triple* mutants. **a-c**, Leaf pieces of Col-0 and (**a**) *pbl30 pbl31 pbl32*, (**b**) *pad4* or (**c**) *adr1 triple* were treated with water (mock) or 500 nM of the indicated elicitor. Boxes indicate total reactive oxygen species (ROS) accumulation (relative light units, RLU) over 30 min (n=16). Data for (**a-c**) corresponds to Fig. 1b, 2b and 3b, respectively. Asterisks indicate results of statistical tests for differences between the mutant and Col-0 response for the given elicitor (Dunnett’s test: ***, p<0.0001; **, p<0.01; *, p<0.05).

**Fig. S6.**
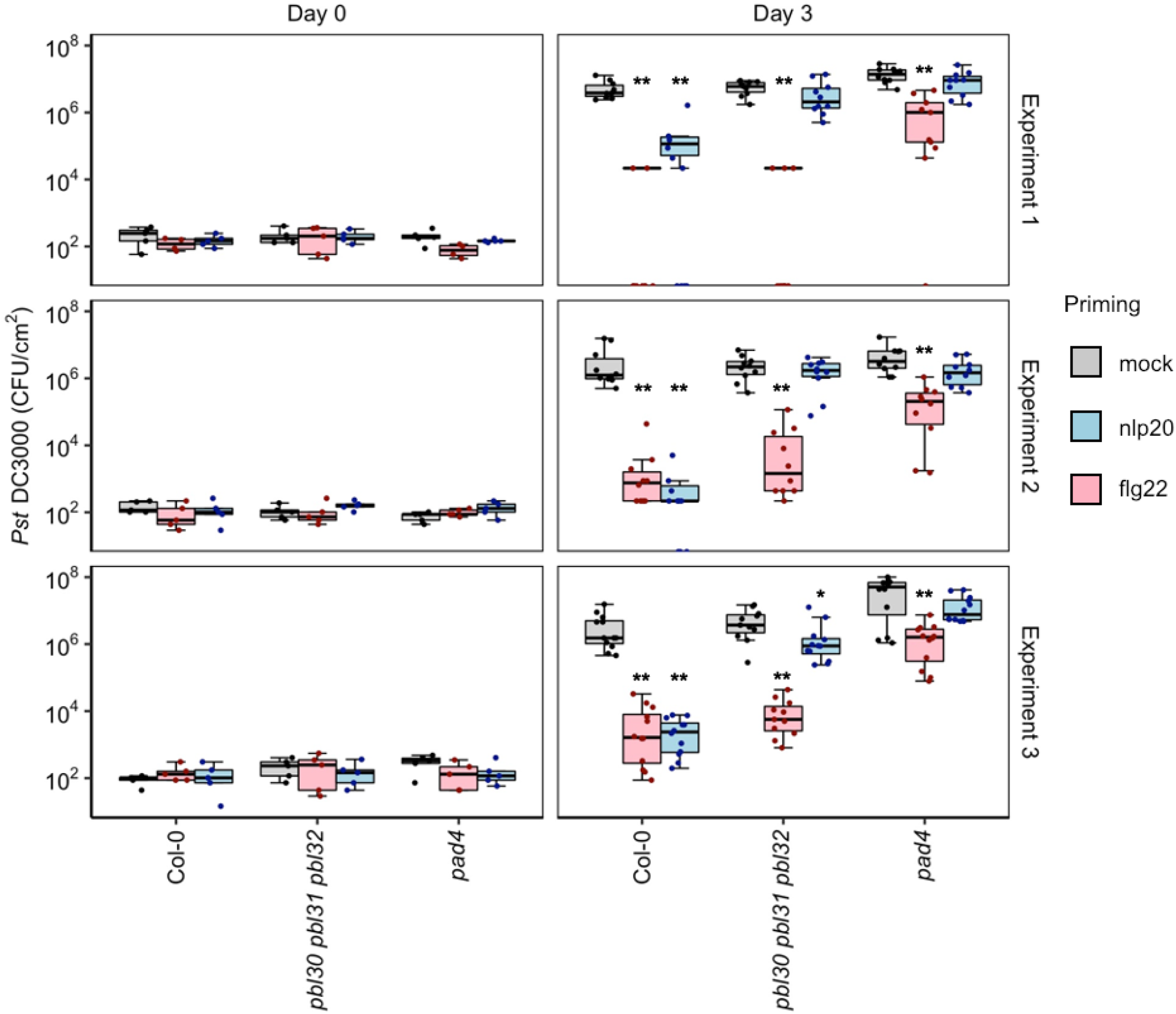
RLCK-VII-7 kinases are required for LRR-RP-mediated priming of enhanced immunity against virulent *Pst* DC3000. Col-0, *pbl30 pbl31 pbl32* and *pad4* leaves were infiltrated with 10 mM MgCl_2_ (mock, grey), 1 μM nlp20 (blue) or 1 μM flg22 (pink). After 24 h, the plants were infiltrated with 10^4^ CFU/mL *Pst DC3000.* Bacterial growth was monitored at day 0 and day 3 (n≥5 for day 0, n≥10 for day 3). Asterisks indicate results of statistical tests for differences between elicitor-primed and mock-treated samples for the indicated plant genotype (Steel’s test: **, p<0.01; *, p<0.05).

**Fig. S7.**
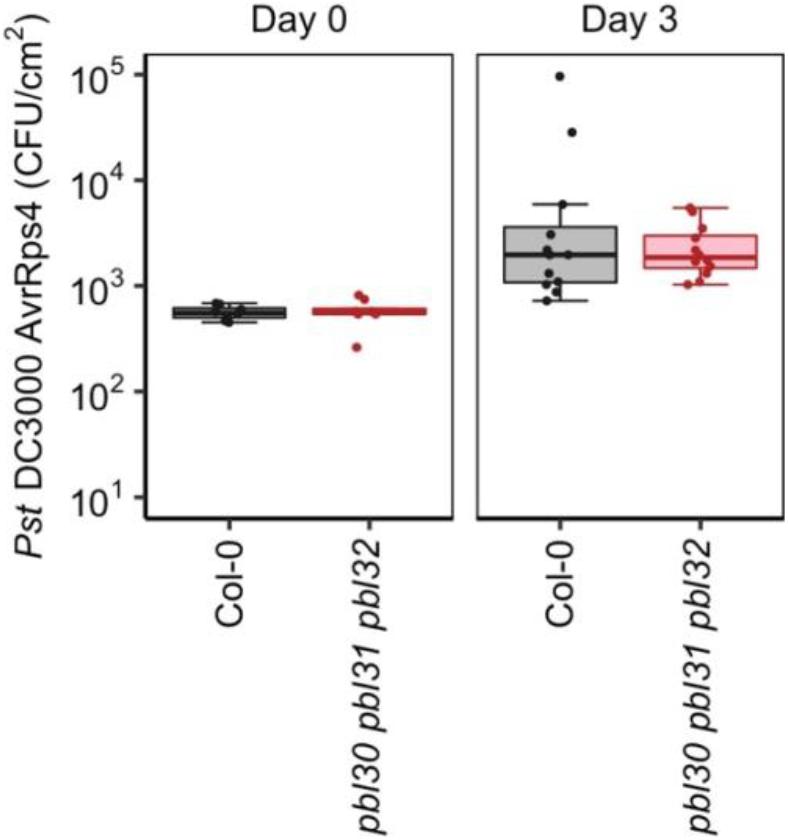
RLCK-VII-7 kinases are not required for an ETI response to *Pst* DC3000 *AvrRPS4*. Col-0 (grey) and *pbl30 pbl31 pbl32* (red) leaves were infiltrated with 10^5^ CFU/mL *Pst* DC3000 *AvrRPS4.* Bacterial growth was monitored at day 0 and day 3 (n=8 for day 0, n=12 for day 3). Steel’s test did not indicate statistically significant differences.

**Fig. S8.**
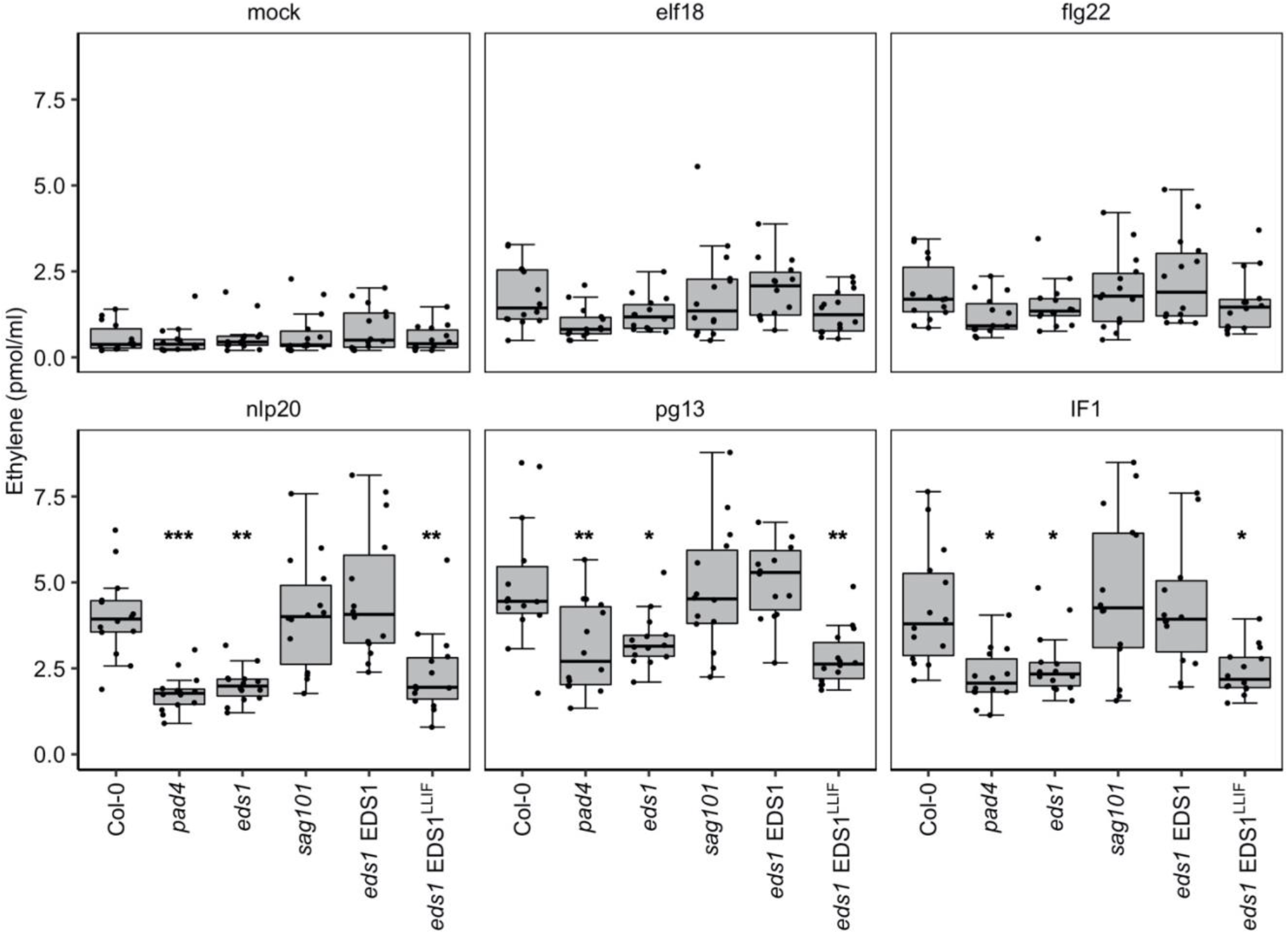
Ethylene accumulation is impaired in *pad4* and *eds1* mutant lines. Leaf pieces of the indicated lines were treated with water (mock), 500 nM elf18, 500 nM flg22, 500 nM nlp20, 500 nM pg13 or 100 nM IF1. Ethylene accumulation was measured after 4 h. Data from three independent experiments (n=13) shown as box plots. Asterisks indicate results of tests for statistical differences (Dunnett’s test vs Col-0: ***, p<0.0001; **, p<0.01; *, p<0.05).

**Fig. S9.**
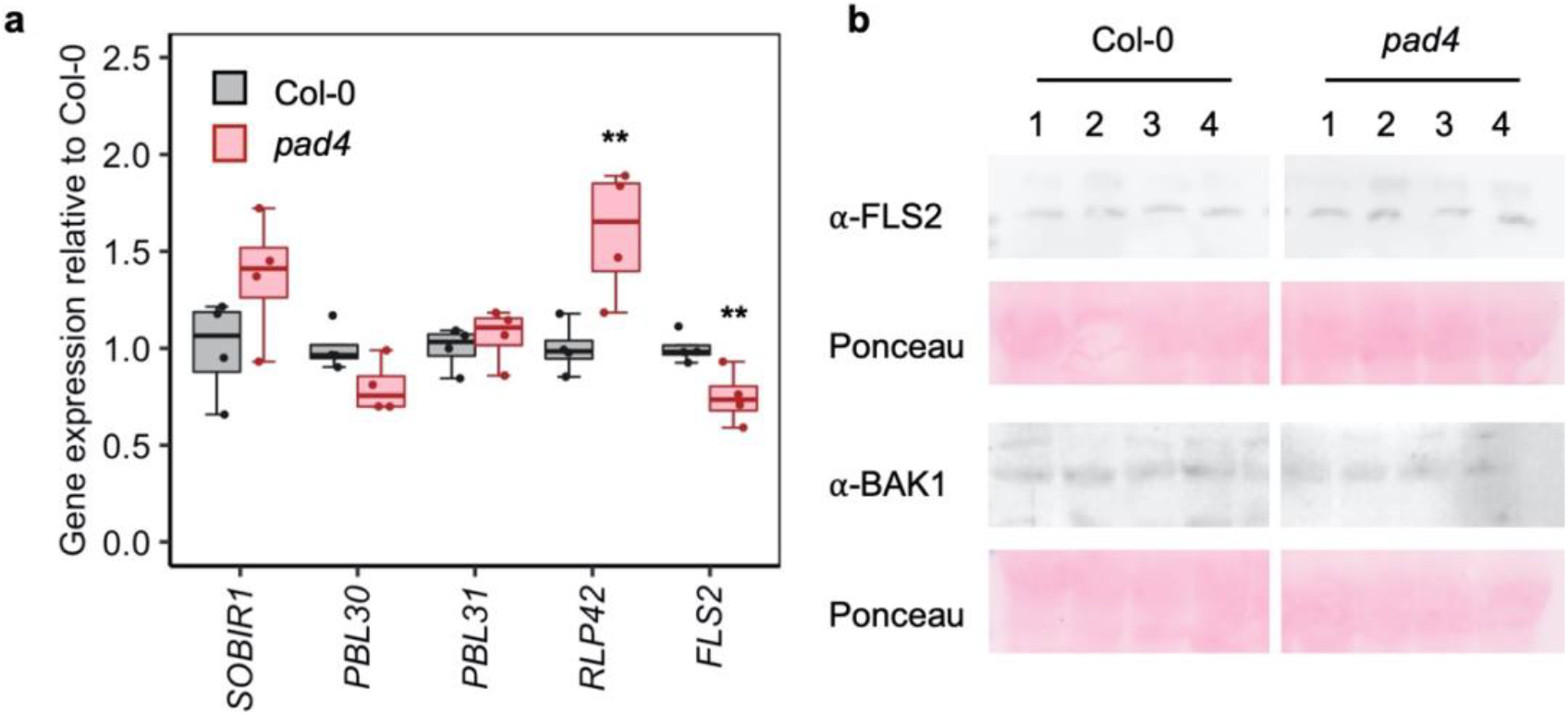
Background levels of immune-related genes are similar in Col-0 and *pad4.* **a**, Transcriptional profiling of *SOBIR1*, *PBL30*, *PBL31*, *RLP42* and *FLS2* by quantitative qRT-PCR. Relative expression of the indicated genes was normalized to the levels of the *EF-1α* transcript and standardized to the levels in Col-0 samples. Data represent one biological experiment with 4 technical replicates. Asterisks indicate results of statistical tests (Dunnett’s test vs Col-0: **, p<0.01). **b**, Protein levels of FLS2 and BAK1 are similar in Col-0 and *pad4.* Two leaves were taken from four 6-week old plants (labeled 1-4) and endogenous BAK1 and FLS2 levels were evaluated by protein blot.

**Fig. S10.**
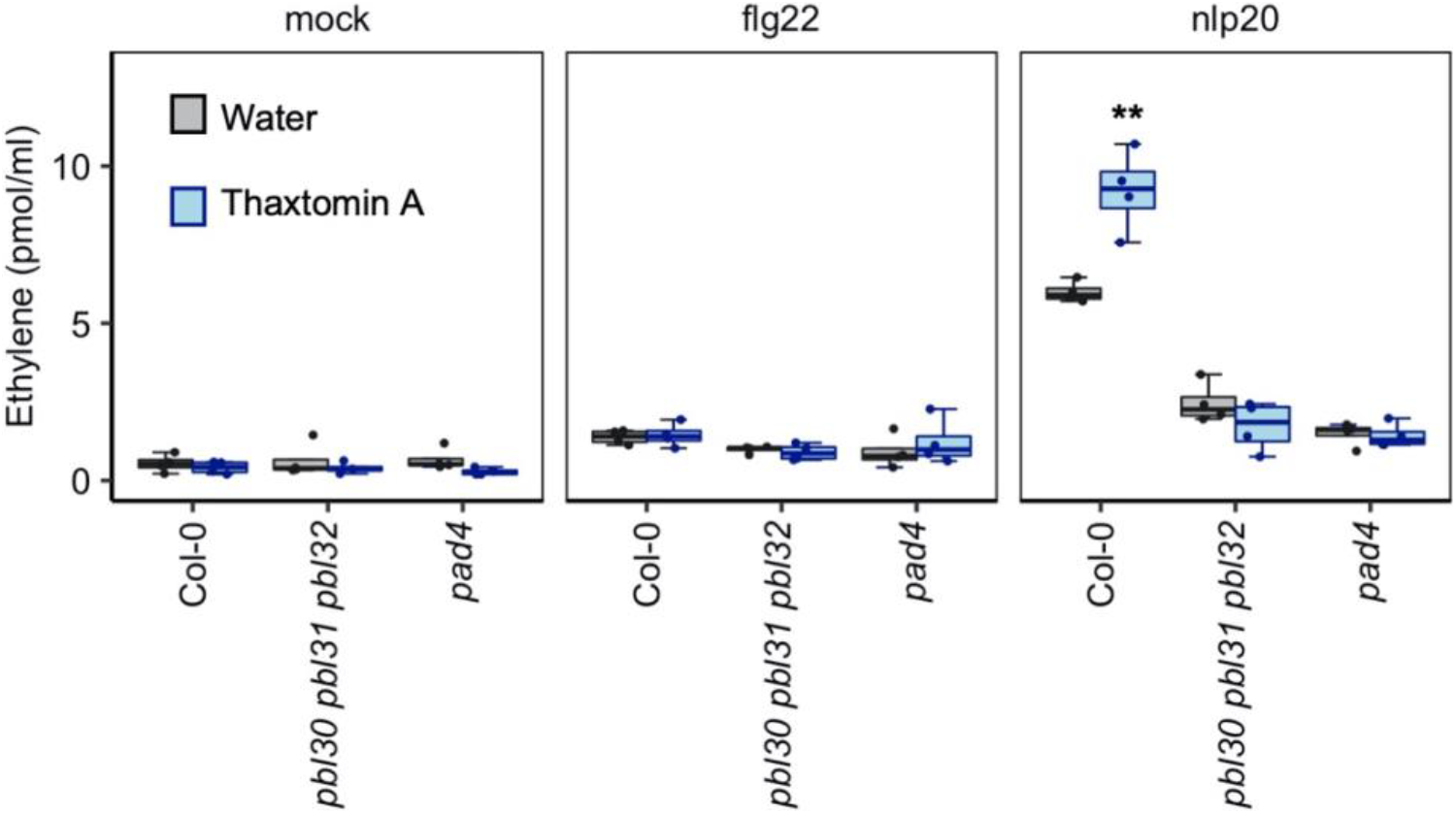
Thaxtomin A (TA) pretreatment enhances nlp20-induced ethylene responses. Leaf pieces of Col-0, *pad4* and *pbl30 pbl31 pbl32* were cut and floated on water (mock, grey) or 100 nM TA (blue) overnight. The next day, the leaf pieces were treated with water (mock), 500 nM nlp20 or 500 nM flg22. Ethylene accumulation was measured after 4 h (n=4). The experiment was repeated with similar results. Asterisks indicate results of statistical tests for differences between TA treated samples compared to the respective water-floated samples (Dunnett’s test: **, p<0.01).

**Fig. S11.**
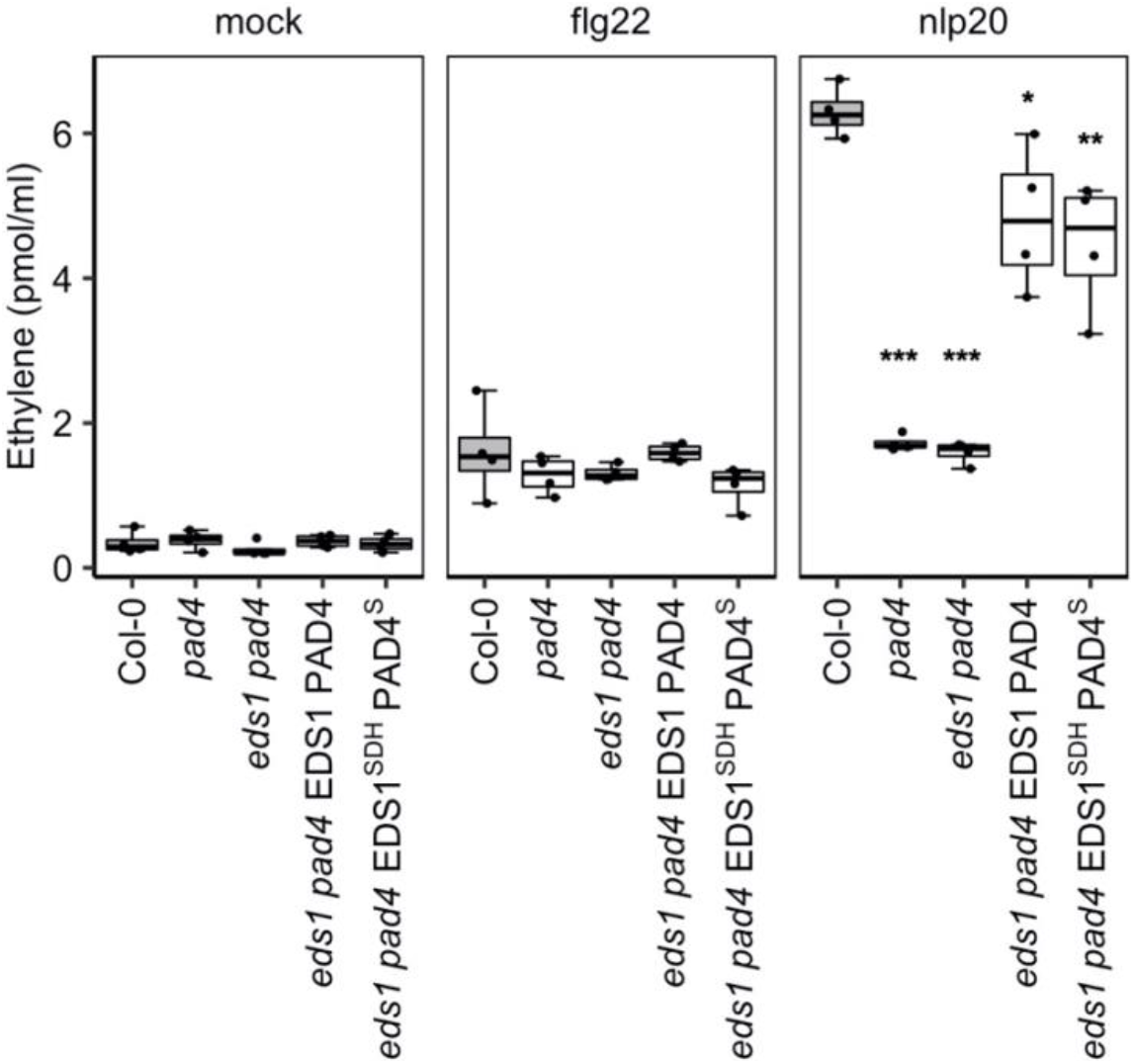
Nlp20 induced ethylene response is not dependent on the PAD4 and EDS1 catalytic residues. Leaf pieces of the indicated lines were treated with water (mock), 500 nM nlp20 or 500 nM flg22. Ethylene accumulation was measured after 4 h (n=4). EDS1^SDH^ and PAD4^S^ harbor mutations in the putative catalytic triads of the two proteins. Note that the putative catalytic residues of PAD4 and EDS1 are not essential for nlp20-induced ethylene response. The experiment was performed three times with similar results. Asterisks indicate results of statistical tests (Dunnett’s test vs Col-0: ***, p<0.0001; **, p<0.01; *, p<0.05).

**Fig. S12.**
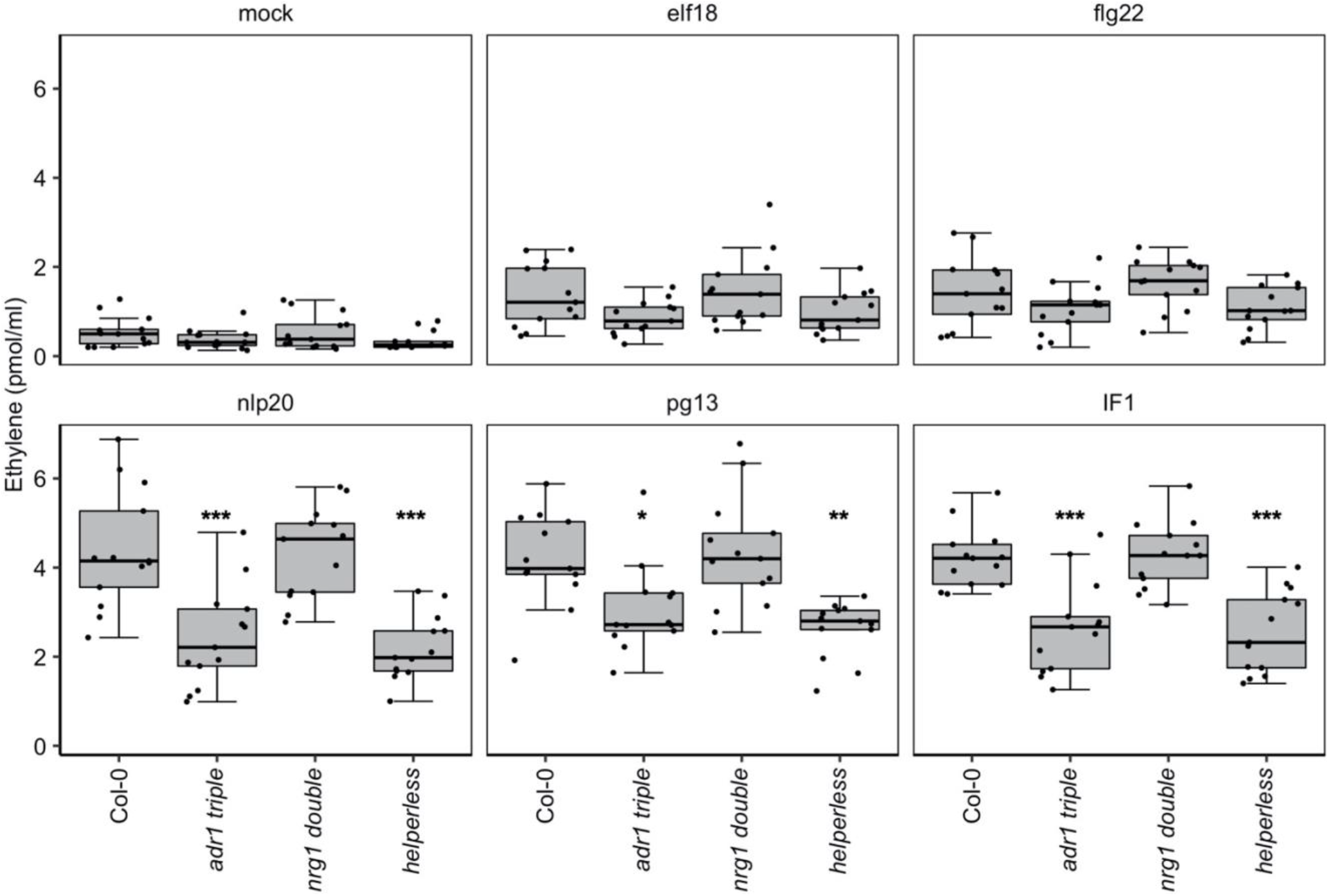
ADR1 helper NLRs are positive regulators of LRR-RP signaling. Leaf pieces of the indicated lines were treated with water (mock), 500 nM elf18, 500 nM flg22, 500 nM nlp20, 500 nM pg13 or 100 nM IF1. Ethylene accumulation was measured after 4 h. Data from three independent experiments shown as box plots with individual datapoints shown as points (n=13). Asterisks indicate results of tests for statistical differences (Dunnett’s test vs Col-0: ***, p<0.0001; **, p<0.01; *, p<0.05).

**Fig. S13.**
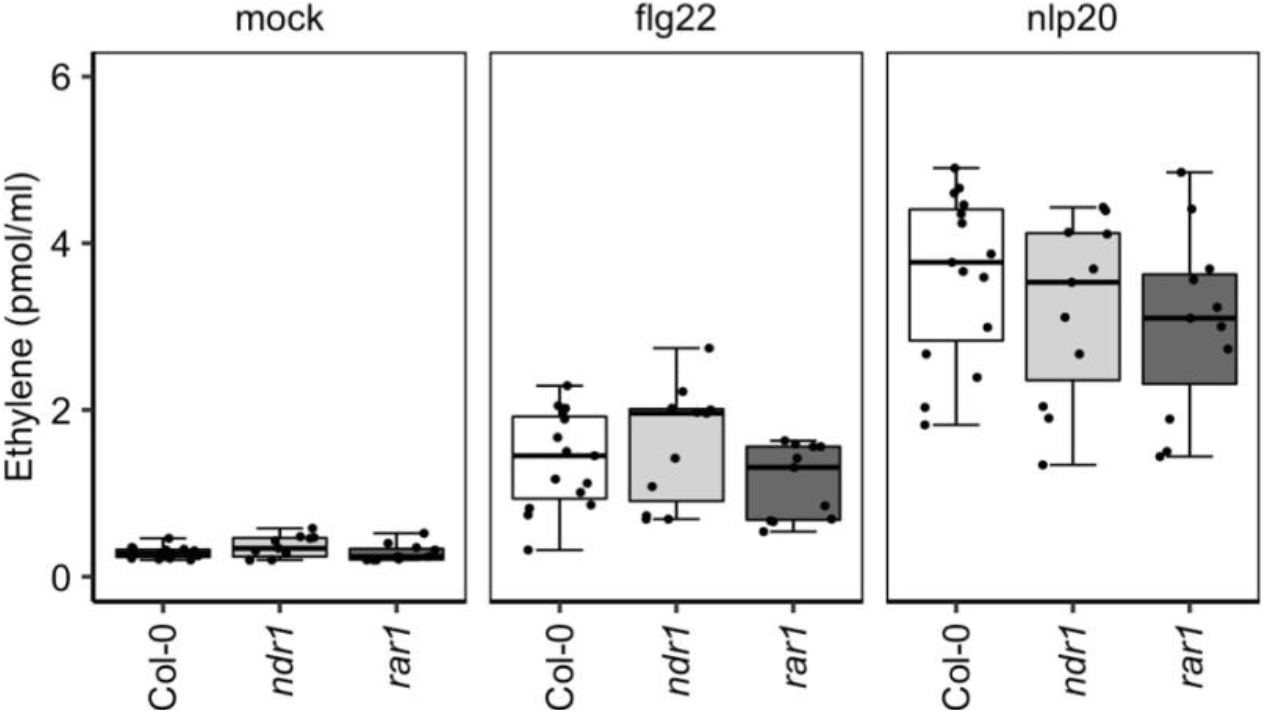
Ethylene responses to nlp20 and flg22 are not impaired in *rar1* or *ndr1* mutant lines. Leaf pieces of Col-0 (white), *ndr1* (light grey) and *rar1* (dark grey) were treated with water (mock) or 500 nM of the indicated elicitor. Ethylene accumulation was measured after 4 h. Composite data from three independent experiments (n≥10) shown as box plots with individual datapoints shown as points. Dunnett’s test did not indicate statistically significant differences between Col-0 and mutant lines.

**Fig. S14.**
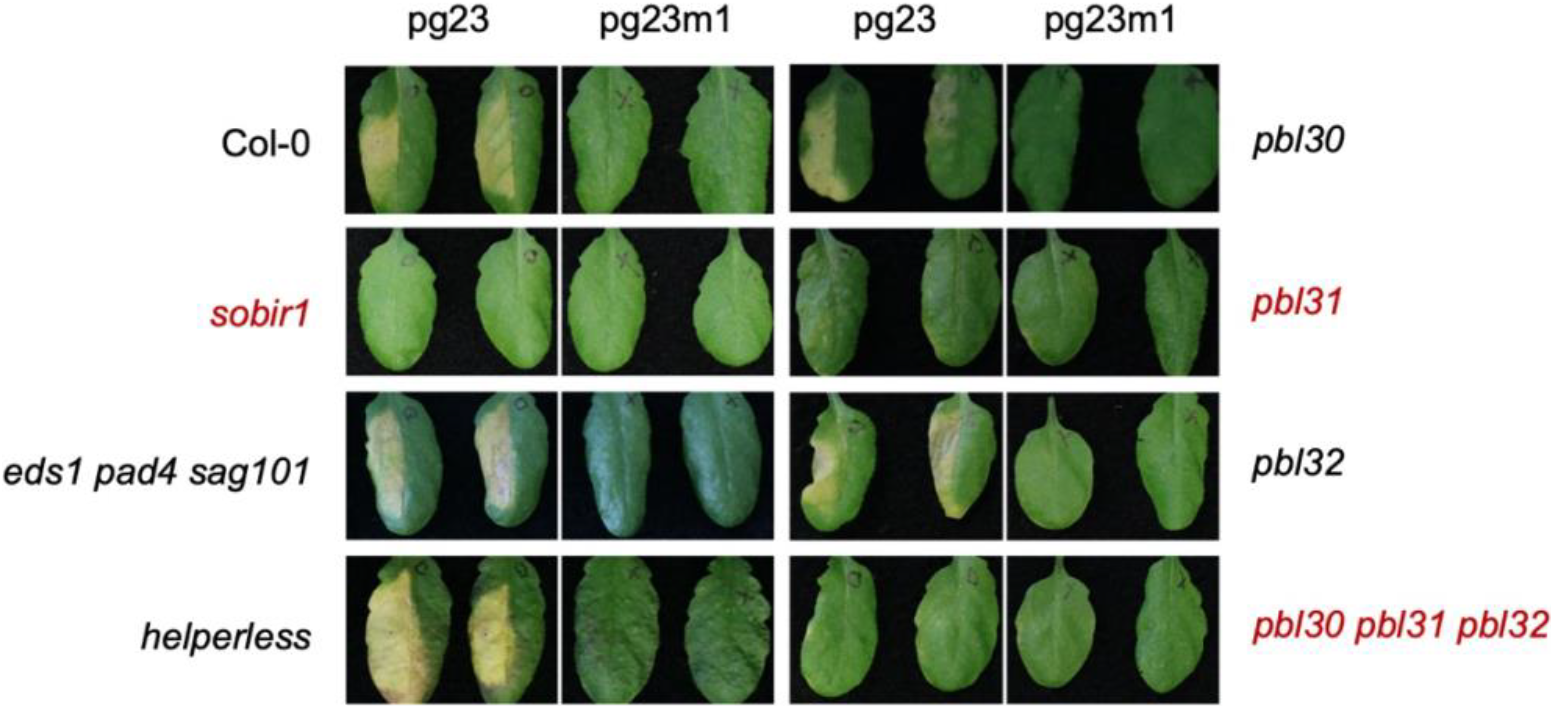
PG-triggered cell death requires SOBIR1 and the RLCK-VII-7 kinase PBL31. *Arabidopsis* leaves were infiltrated with 10 μM pg23 or the inactive variant pg23m1. Chlorosis and lesion formation were visible after 7 days. Lines without visible cell death upon pg23 infiltration are marked in red.

**Fig. S15.**
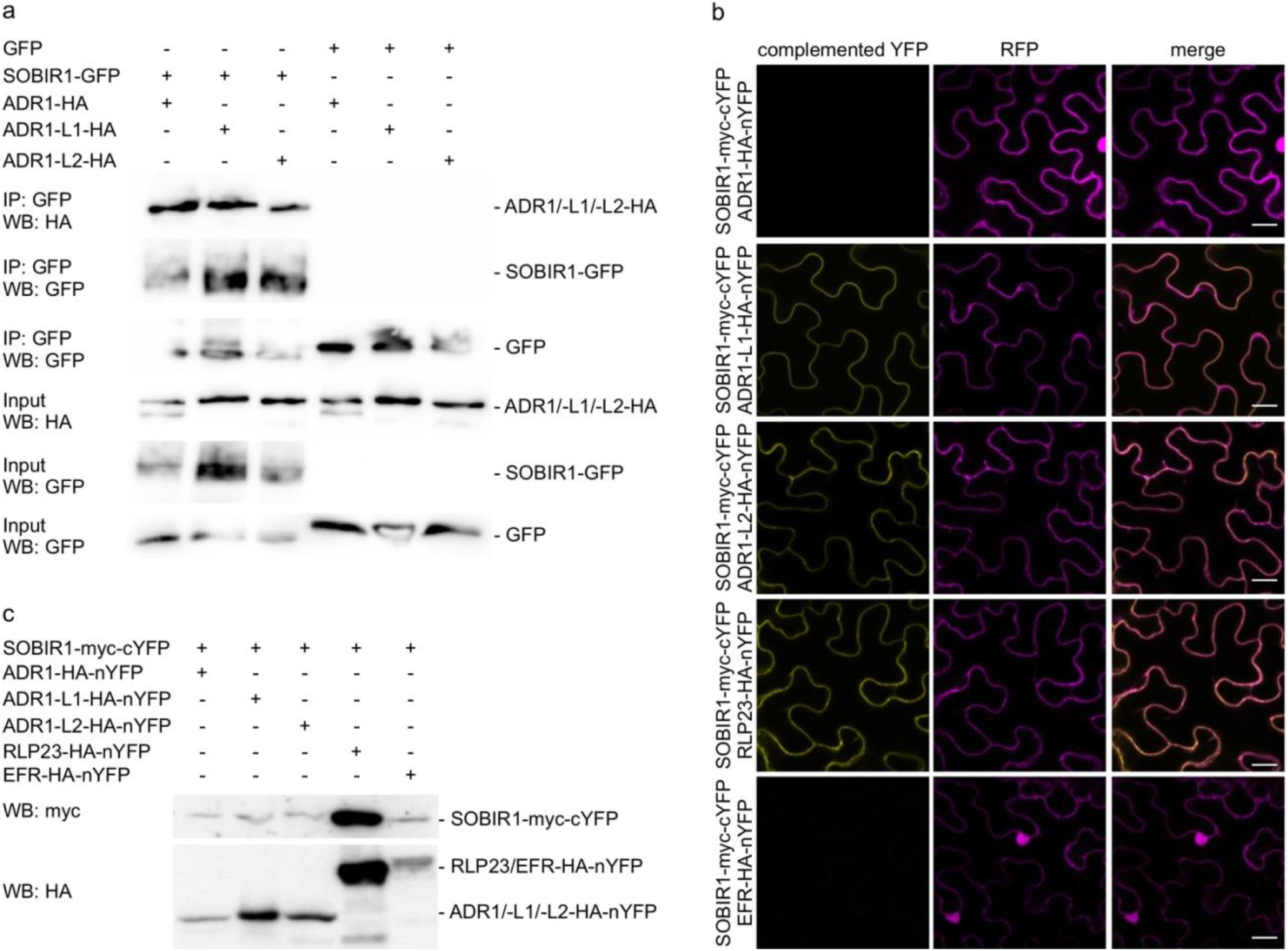
ADR1s associate with SOBIR1. **a**, Pull-down of GFP and SOBIR1-GFP transiently co-expressed with ADR1-HA, ADR1-L1-HA or ADR1-L2-HA. Plants transiently expressing the different proteins were subjected to co-immunoprecipitation using GFP-trap beads and subsequently analyzed by protein blot using tag-specific antisera. **b**, BiFC between SOBIR1 and the ADR1s confirms constitutive interaction of SOBIR1 with ADR1-L1 and ADR1-L2 at the plasma membrane. **c**, Protein levels of the transiently expressed proteins in BiFC experiments shown in panel (**b**).

## Notes

### Competing Interest Statement

The authors have declared no competing interest.

